# Mechanistic characterization of *RASGRP1* variants identifies an hnRNP K-regulated transcriptional enhancer contributing to SLE susceptibility

**DOI:** 10.1101/568790

**Authors:** Julio E. Molineros, Bhupinder Singh, Chikashi Terao, Yukinori Okada, Jakub Kaplan, Barbara McDaniel, Shuji Akizuki, Celi Sun, Carol Webb, Loren L. Looger, Swapan K. Nath

## Abstract

Systemic lupus erythematosus (SLE) is an autoimmune disease with a strong genetic component. We recently identified a novel SLE susceptibility locus near *RASGRP1*, which governs the ERK/MAPK kinase cascade and B-/T-cell differentiation and development. However, precise causal *RASGRP1* functional variant(s) and their mechanisms of action in SLE pathogenesis remain undefined. Our goal was to fine-map this locus, prioritize genetic variants according to likely functionality, experimentally validate the contribution of three SNPs to SLE risk, and experimentally determine their biochemical mechanisms of action. We performed a meta-analysis across six Asian and European cohorts (9,529 cases; 22,462 controls), followed by *in silico* bioinformatic and epigenetic analyses to prioritize potentially functional SNPs. We experimentally validated the functional significance and mechanism of action of three SNPs in cultured T-cells. Meta-analysis identified 18 genome-wide significant (p<5×10^−8^) SNPs, mostly concentrated in two haplotype blocks, one intronic and the other intergenic. Epigenetic fine-mapping, allelic, eQTL and imbalance analyses predicted three transcriptional regulatory regions with four SNPs (rs7170151, rs11631591-rs7173565, and rs9920715) prioritized for functional validation. Luciferase reporter assays indicated significant allele-specific enhancer activity for intronic rs7170151 and rs11631591-rs7173565 in T-lymphoid (Jurkat) cells, but not in HEK293 cells. Following up with EMSA, mass spectrometry and ChIP-qPCR, we detected allele-dependent interactions between heterogeneous nuclear ribonucleoprotein K (hnRNP-K) and rs11631591. Furthermore, inhibition of hnRNP-K in Jurkat and primary T-cells downregulated *RASGRP1* and ERK/MAPK signaling. Comprehensive association, bioinformatics, and epigenetic analyses yielded putative functional variants of *RASGRP1*, which were experimentally validated. Notably, intronic variant (rs11631591) is located in a cell type-specific enhancer sequence, where its risk allele binds to the hnRNP-K protein and modulates *RASGRP1* expression in Jurkat and primary T-cells. As risk allele dosage of rs11631591 correlates with increased *RASGRP1* expression and ERK activity, we suggest that this SNP may underlie SLE risk at this locus.

## Introduction

Systemic lupus erythematosus (SLE) is a complex autoimmune disease that disproportionately affects people of Asian, African, and Hispanic ethnicities and women, in particular, with higher incidence and disease severity [1]. Much of SLE etiology remains mysterious. It has been proposed that complex interactions amongst numerous genes and their products with pathogens and other environmental factors promotes dysregulation of both the innate and adaptive immune responses in SLE. Over 80 SLE susceptibility loci have been identified so far across multiple ethnic groups by genome-wide association studies (GWAS) and candidate gene studies [2, 3]. However, the precise underlying variants and functional mechanisms associated with disease are largely unidentified for the vast majority of these SLE-associated signals. Understanding SLE pathogenesis requires identification of true causal variants and the target genes and mechanisms by which they contribute to disease.

Previously, we reported a novel SLE susceptibility signal near the RAS guanyl-releasing protein 1 (*RASGRP1*) in Asians [4]. We identified several associated variants, the most significant being an intergenic variant (rs12900339) between *RASGRP1* and *C15orf53* [4]. However, the actual predisposing variants, target genes, and underlying mechanisms of action for this region are largely unknown. *RASGRP1* belongs to a family of RAS guanyl nucleotide-releasing proteins (RASGRPs) comprising four members (*RASGRP1* through *RASGRP4*), all with a diacylglycerol (DAG)-binding C1 catalytic domain. Upon antigen stimulation, DAG binding and phospholipase C (PLC) signaling drive RASGRPs to the membrane, where they play important roles in RAS activation [5, 6]. *RASGRP1*, originally cloned from the brain [7], was later found highly expressed in T-lymphocytes [8]; small amounts of *RASGRP1* expression can also occur in B-lymphocytes, neutrophils, mast cells, and natural killer cells [9-11]. *RASGRP1* has been shown to be involved in B-cell development, activation and tolerance, in both mice and humans[12, 13]. *RASGRP1*^−/−^ mice have been reported for marked deficiency in thymocyte and lymphocyte development, which was associated with impaired proliferation in response to TCR stimulation[14]. Deficiency in *RASGRP1* in mice, has been associated with CD4+ and CD8+ T cell lymphopenia[8]. However, in humans deficient in *RASGRP1* show decrease in CD4+T concurrent with a relative increase in CD8+T cells[15]. *RASGRP1* inhibition impairs T-cell expansion and increases susceptibility to Epstein-Barr virus infection, as well as suppressing proliferation of activated T-cells occurring in autoimmune conditions [16]. A recent study reported a heterozygous mutation in *RASGRP1* correlated with autoimmune lymphoproliferative syndrome (ALPS)-like disease[17]. *RASGRP1* expression in T-cells also correlated negatively with rheumatoid arthritis disease activity[18]. Dysregulated expression of *RASGRP1* has been observed in human SLE. The ratio of normal *RASGRP1* isoforms to isoforms missing exon-11 could be linked to defective poly[ADP-ribose] polymerase 1 (*PARP1*) expression and reduced lymphocyte survival in SLE patients [19, 20]. Aberrant splice variants accumulate in SLE patients and adversely affect T-cell function[21]. There are conflicting reports of the effect of *RASGRP1* on ERK signaling. In one hand, deficiency in *RASGRP1* expression reportedly decreases ERK phosphorylation in B- and T-cells[15]. Hydralazine, a drug that causes drug-induced lupus erythematosus, is reported to inhibit ERK signaling, inducing autoimmunity and the production of anti-dsDNA autoantibodies in mice [22]. However, some reports found significantly higher levels of pERK and pJNK in SLE patients with active disease *versus* controls and inactive SLE patients[23-25], contradicting earlier reports. In spite of these conflicting reports, the consensus is that *RASGRP1* dysfunction is mechanistically associated with autoimmune phenotypes including SLE.

Here, we fine-mapped an SLE locus near *RASGRP1* that we previously identified [4]. Using trans-ethnic meta-analysis across six Asian and European cohorts followed by bioinformatic analyses and experimental validation, we identified potential SLE predisposing variants and defined mechanisms by which these functional variants contribute to SLE pathogenicity.

### Materials and Methods

#### Patients and Data

We used all associated SNP data at this locus from six cohorts reported previously (**Table 1**). We began with our published Asian cohort report **(see Supplementary Table 5 in** [4]) and augmented this with two publicly available sets of GWAS summary statistics [26, 27] and a partially published Japanese cohort [28]. Our original report contained three Asian cohorts (Korean, Han Chinese, and Malaya Chinese). Japanese samples included samples (456 cases and 1,102 controls) collected under support of the Autoimmune Disease Study Group of Research in Intractable Diseases, Japanese Ministry of Health, Labor and Welfare, and the BioBank Japan Project [28], and added samples obtained at Kyoto University, Japan. SLE classification followed the American College of Rheumatology criteria [29]. All sample collections were approved by the Institutional Review Board of the Oklahoma Medical Research Foundation as well as by the collaborating institutions.

**Table 1.** Cohorts used in this study. We utilized samples from our previous report Sun *et al.* 2016 for *RASGRP1* SLE association. We added a Han Chinese (HC) and a European (EU) cohort from Morris *et al.* 2016 and a Japanese cohort containing the patients from Okada *et al.* 2012 and additional Japanese samples (JAP).

| Population | Cohort | Cases | Controls | Publication |
| --- | --- | --- | --- | --- |
| Asian | Korean | 2,487 | 7,620 | Sun et al 2016 |
|  | Han Chinese |  |  |  |
|  | Malaya Chinese |  |  |  |
| HC | Han Chinese | 1,659 | 3,398 | Morris et al 2016 |
| EU | European | 4,036 | 6,958 | Bentham et al 2015 |
| JAP | Japanese | 1,347 | 4,486 | Okada et al. 2012 +<br>new Data |
| TOTAL |  | 9,529 | 22,462 |  |

#### Quality Control

SNP quality control for our initial Asian cohort has been described elsewhere [4]. Quality control for European, Han Chinese 2, and Japanese samples was described in the original publications [26-28]. All SNPs in the study were in Hardy-Weinberg equilibrium (P>1×10^−6^), and had minor allele frequency >0.5%. Genotypic missingness was <10%. In order to match risk alleles between cohorts, we compared their allele frequencies to the parent populations from the 1000 Genomes Project. We used the SNP reference dbSNP142 as the SNP-naming convention in common for all variants. SNP imputation for all cohorts was described in their original publications. For this study, SNPs with r^2^ and imputation quality information <0.7 were dropped.

#### Study design

In order to identify *RASGRP1* functional variants and their mechanisms of action, our analysis followed the workflow presented in **Figure 1**. We first extracted all summary GWAS information in and around *RASGRP1* (118 SNPs) from Supplementary Table 5 in our previous study of Asian SLE [4]. We combined these results with a European [26], an Asian [27], and a partially published Japanese cohort [28], to perform meta-analysis. SNPs that passed the genome-wide significant association threshold (p = 5×10^−8^) were further annotated with functional information. A series of bioinformatics and epigenomic analyses was conducted for each of the candidate SNPs including their effects on gene expression (expression quantitative trait loci, eQTLs), transcription factor binding, promoter/enhancer activities and chromatin interaction sites. Together, we prioritized and nominated SNPs with stronger association signals and with higher annotated likelihood of being functional (**Supplementary Table 1 and 2**). Finally, we experimentally validated predicted functions of the nominated SNPs in Jurkat and HEK293 cell lines. Following SNP prioritization, we performed electrophoretic mobility shift assays (EMSAs), followed by mass spectrometry, chromatin immuno-precipitation quantitative PCR (ChIP-qPCR), and inhibition-based expression assays.

**Table 2.**
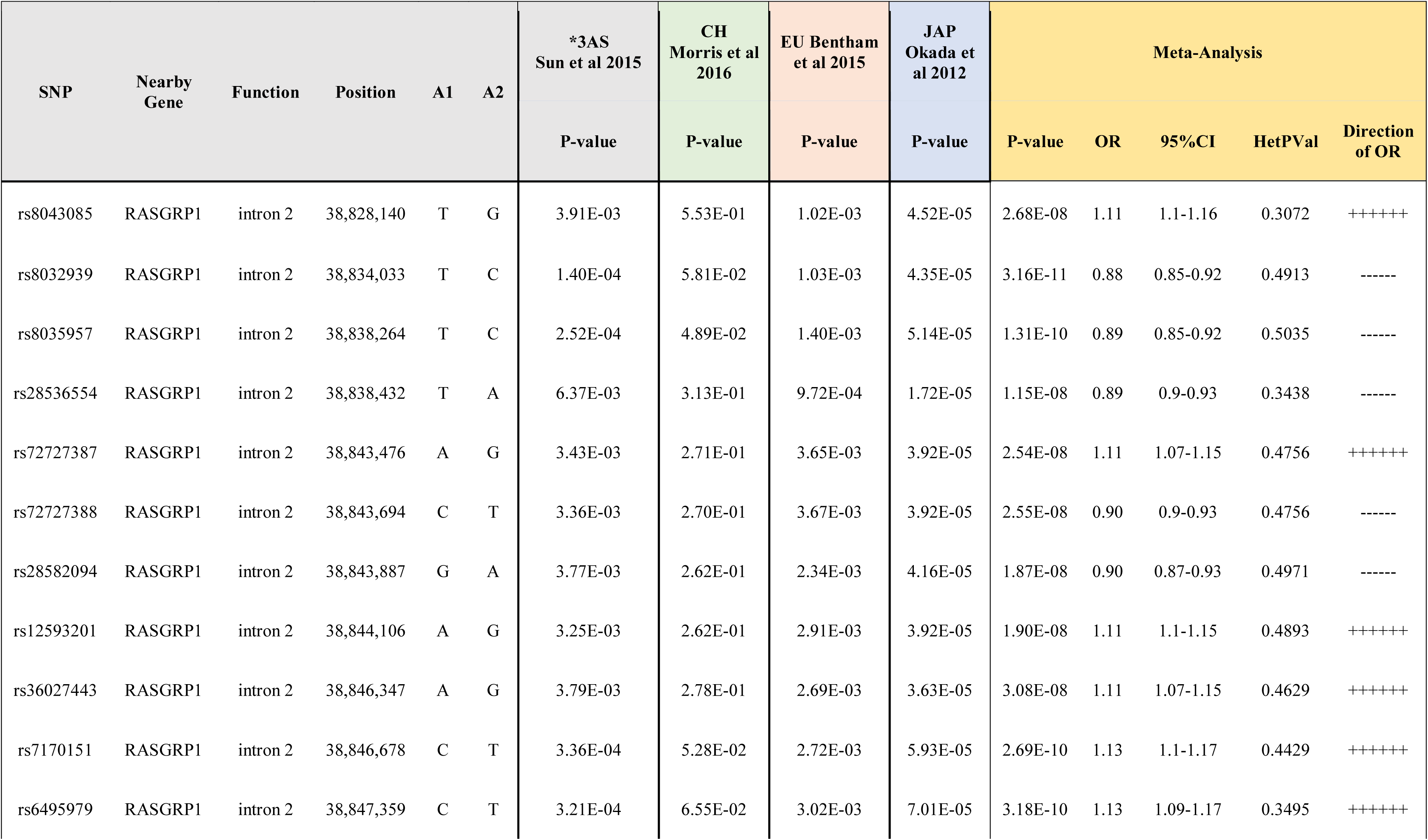
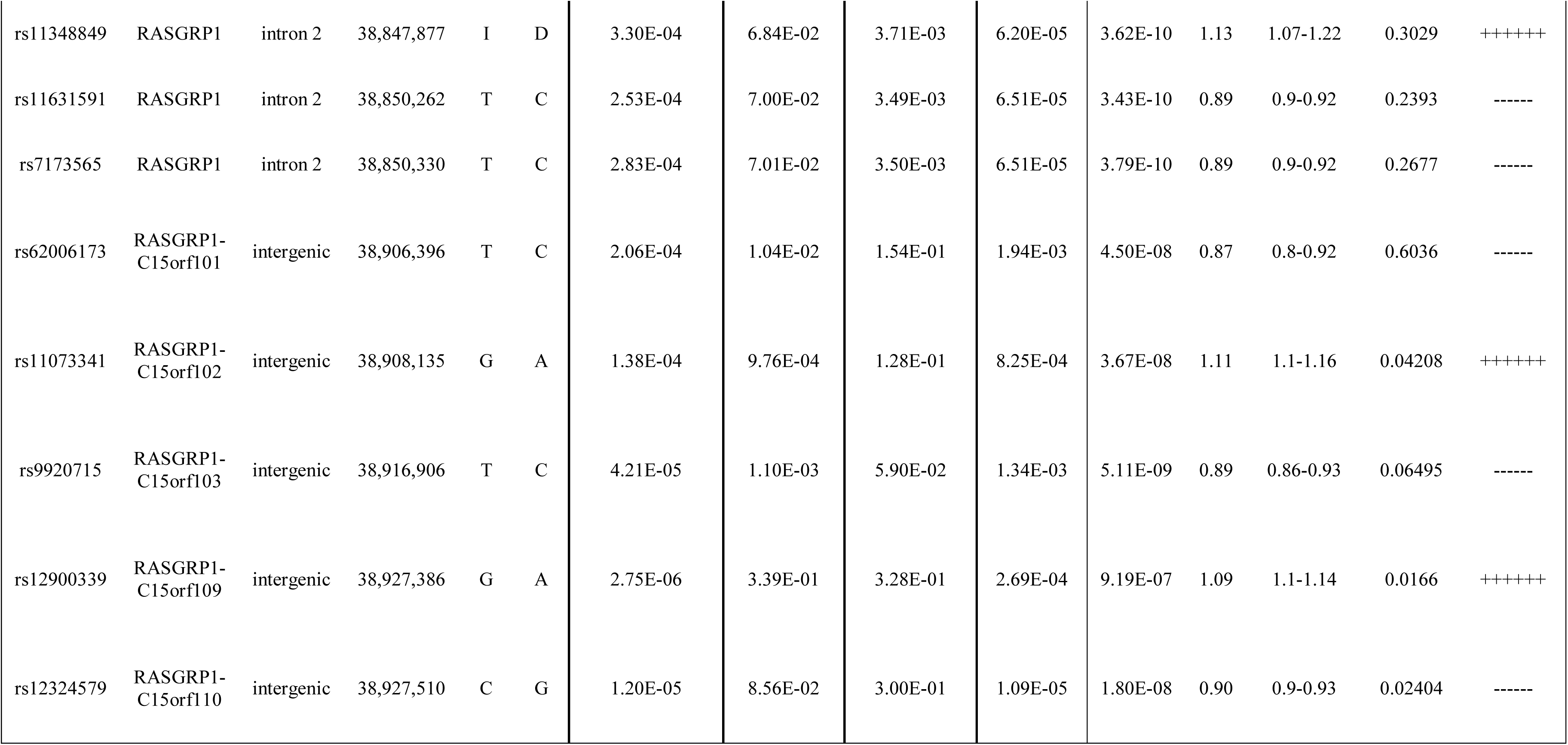
Meta-analysis of the *RASGRP1* region. We identified 18 genome-wide significant SNPs in *RASGRP1* (intron 2) and between *RASGRP1* and *C15orf53* (intergenic). *Sun *et al.* 2016 supplementary table 5 (Korean, Han-Chinese, Malay Chinese, 3AS) was used as the discovery cohort and was replicated in Morris *et al.* 2016 (Han Chinese, HC); and Bentham *et. al.* 2015 (European, EU) and in Okada *et al.* 2012 + new samples (Japanese, JAP) cohorts. OR: Odds ratio. 95% CI: 95% confidence interval. HetPVal: P-value for heterogeneity meta-analysis test. Direction of OR is presented as + if OR >1 and – if OR<1. Note that all OR directions are consistent for all SNPs.

**Figure 1.**
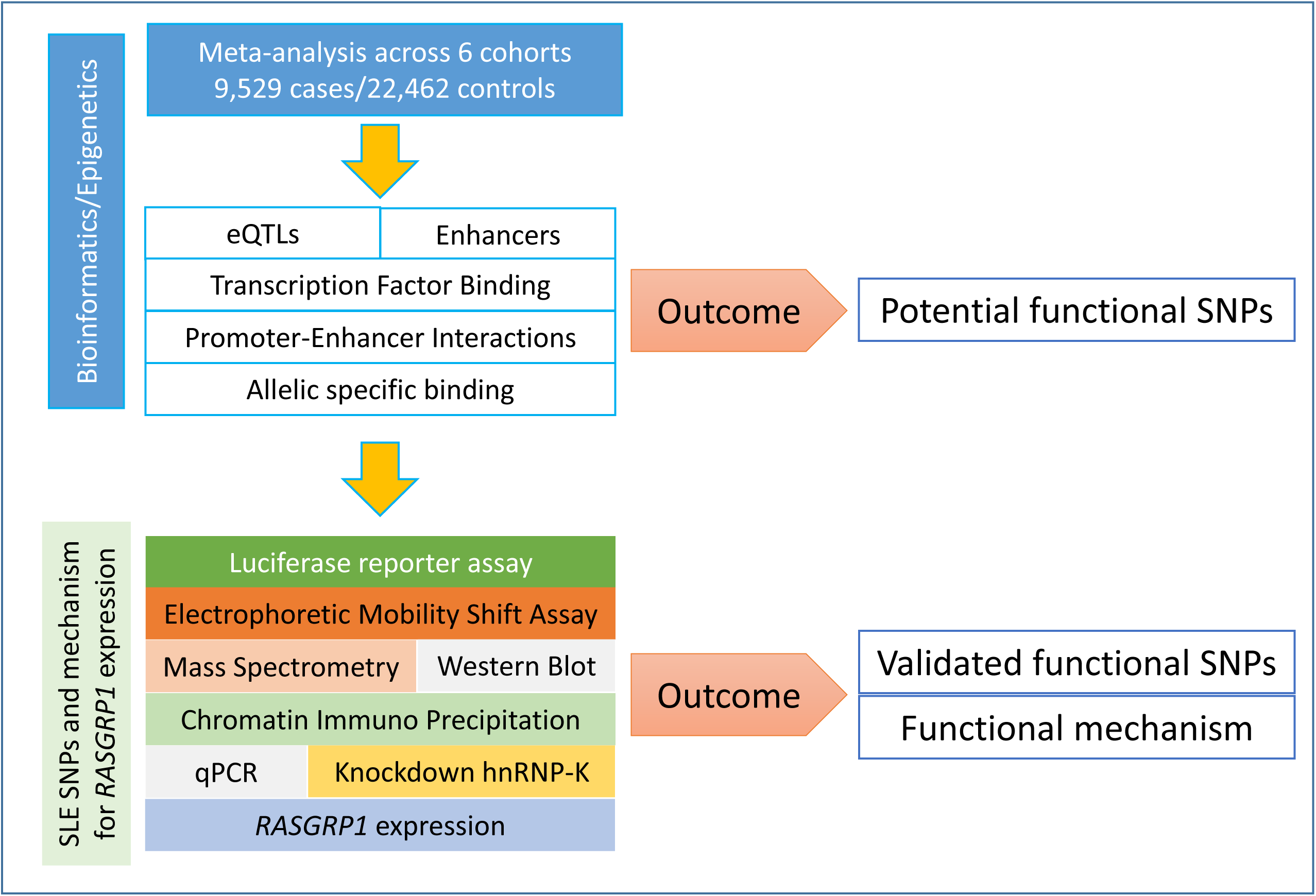
Framework of study design. Our study followed our bioinformatics-prioritized potential functional SNPs with laboratory validation along many different dimensions.

#### Association analysis and trans-ethnic meta-analysis

Association analysis for all cohorts was performed using PLINK [30] and SNPTEST. Meta-analysis for all cohorts was performed in METAL [31] using cohort sample size correction and standard error correction to estimate the 95% confidence interval for odds ratios. Heterogeneity of odds ratios was estimated and informed the use of P_meta_ values in the study. Variants with P_meta_<5×10^−3^ were selected for further study.

#### Bioinformatic analysis

Given that candidate SNPs were located in non-coding regions of the genome, we performed a thorough epigenetic annotation of the variants. Initial annotation of epigenetic features was performed in Haploreg [32]. Each SNP in the region was collocated with active and regulatory histone marks including H3K27ac, H3K4me1 and H3K4me3, and DNase hypersensitivity sites (DHS) in GM12878, and CD4+ and CD8^+^ T cells (**Supplementary Figure 1**). Histone modifications and DHS data were obtained from the ENCODE project [33] and the BLUEPRINT epigenome project [34].

#### SNP prioritization

We used a prioritization algorithm to narrow down the large list of SNPs for further validation. Our strategy consisted of two Bayesian algorithms to score each SNP (3dSNP [35] and RegulomeDB [36]), as well as additional expression, epigenetic, and preferential allele-specific information about each SNP. First we used the 3dSNP [35] tool to assign functional weights based on the presence of enhancers, promoters, experimentally determined (ChIP-seq) transcription factor binding sites (TFBSs), TFBS motif matching, evolutionary conservation, and presence of 3D chromatin interactions. We assigned a 3dSNP weight of 2 to SNPs greater than two standard deviations above the mean, a weight of 1 for scores above the mean, and a weight of 0 for the rest. RegulomeDB [36] scores were also assigned for each candidate SNP, and converted to an associated weight. Each functional category, *i.e.* eQTL, enhancer/super-enhancer, rSNP [37], capture Hi-C, TFBS, and allele-specific expression/binding, was assigned a weight of 1 if the SNP had this feature. Finally, we summed all weights for each SNP and nominated the top SNPs for further experimental validation.

#### Expression quantitative trait loci (eQTLs)

All the candidate SNPs were annotated for the presence of eQTLs changing expression of *RASGRP1* and its surrounding genes in multiple tissues. We used expression databases for whole blood [38, 39], immune cell lines [40], and multiple tissues [41] (GTEx Analysis Release V6p). In order to identify quantitative changes in methylation in blood cell lines, we used the WP10 database from the Blueprint epigenome project [42].

#### Transcription factor binding sites (TFBSs)

In order to identify allele-specific effects on transcription factor binding (TFBSs), we used the motifBreakr [43] algorithm implemented in R, as well as the PERFECTOS-APE algorithm that identifies fold-changes in binding affinity of SNPs against HOCOMOCO10, HOMER, JASPER, Swiss Regulon, and HT-Selex motif databases. We selected only TFBSs that had at least 5-fold change in affinity.

#### Assessing SNP effects on enhancer/promoter sequences

We assessed whether each SNP was located within regulatory (enhancer/promoter) regions across multiple cell lines using active histone marks (H3K27ac, H3K4me1 and H3K4me3) collocation implemented in the 3dSNP application [35]. Super-enhancers were annotated using the dbSuper [44], Prestige [45] and EnhancerAtlas [46] databases.

#### Chromatin interactions

Chromatin looping was identified using capture Hi-C assays obtained from 3D Genome [47], 3DSNP [35] and CHiCP [48]; as well as from Promoter-capture Hi-C [49-52] experiments.

#### Allele-specific binding

Candidate SNPs within the association peaks were further targeted to assess allele-specific binding (ASB) of histone marks H3K4me1 and H3K4me3 in and around them. ASB was calculated using seven heterozygous cell lines (GM10847, GM12890, GM18951, GM19239, GM19240, GM2610, and SNYDER). ASB was implemented in SNPhood [53].

#### Luciferase reporter assays

To test candidate SNP-containing regions for allele-specific enhancer activity, we cloned all three SNPs (rs1163159, rs7173565-rs7173565, and rs9920715) individually into the enhancer reporter plasmid pGL4.26[luc2/minP/Hygro] (Promega, USA). In brief, genomic DNA from the Coriell cell line having different genotypes for the SNP tested (obtained from NIGMS Human Genetic Cell Repository at the Coriell Institute for Medical Research) was amplified using specific primers containing KpnI and HindIII sites (**Supplementary Table 3**). These amplified PCR products surrounding rs11631591 (481 bp), rs7170151 (579 bp) and rs9920715 (455 bp) were digested with KpnI and HindIII restriction enzymes and ligated into the pGL4.26 plasmid. After cloning and transformation, the plasmids generated for each genotype were confirmed by direct Sanger DNA sequencing. To study cell type-dependence, we used two different cell types, human embryonic kidney HEK293 and T-lymphoid Jurkat cell lines. HEK293 cells were seeded in 24-well sterile plastic culture plates at a density of 1×10^5^ cells per well with complete growth medium. The cells were transfected with 500 ng of pGL4.26 (with or without insert) along with 50 ng *Renilla* plasmid as control vector to control for differences in transfection efficiency. LipofectAMINE 3000 (Invitrogen, USA) was used for transfection into HEK293 cells, according to the manufacturer’s protocol. For Jurkat transfections, we used the Neon Transfection System (Thermo Fisher Scientific).□A total of 5 × 10^5^ Jurkat cells was electroporated with a Neon Transfection System (Invitrogen) under the following conditions: voltage (1050 V), width (30 ms), pulses (Two), 10-μl tip, and Buffer R. For transfection, 2□μg of each plasmid containing the insert with risk or non-risk allele, along with 50 ng *Renilla* plasmid. Firefly and *Renilla* luciferase activities were measured consecutively at 24h after transfection using Dual-luciferase assays (Promega) according to the manufacturer’s instructions. Luciferase activity was analyzed with Student’s t-test implemented in GraphPad Prism7. Differences between relative luciferase activity levels were considered significant if Student’s t-test *P*-value <0.05.

**Table 3.**
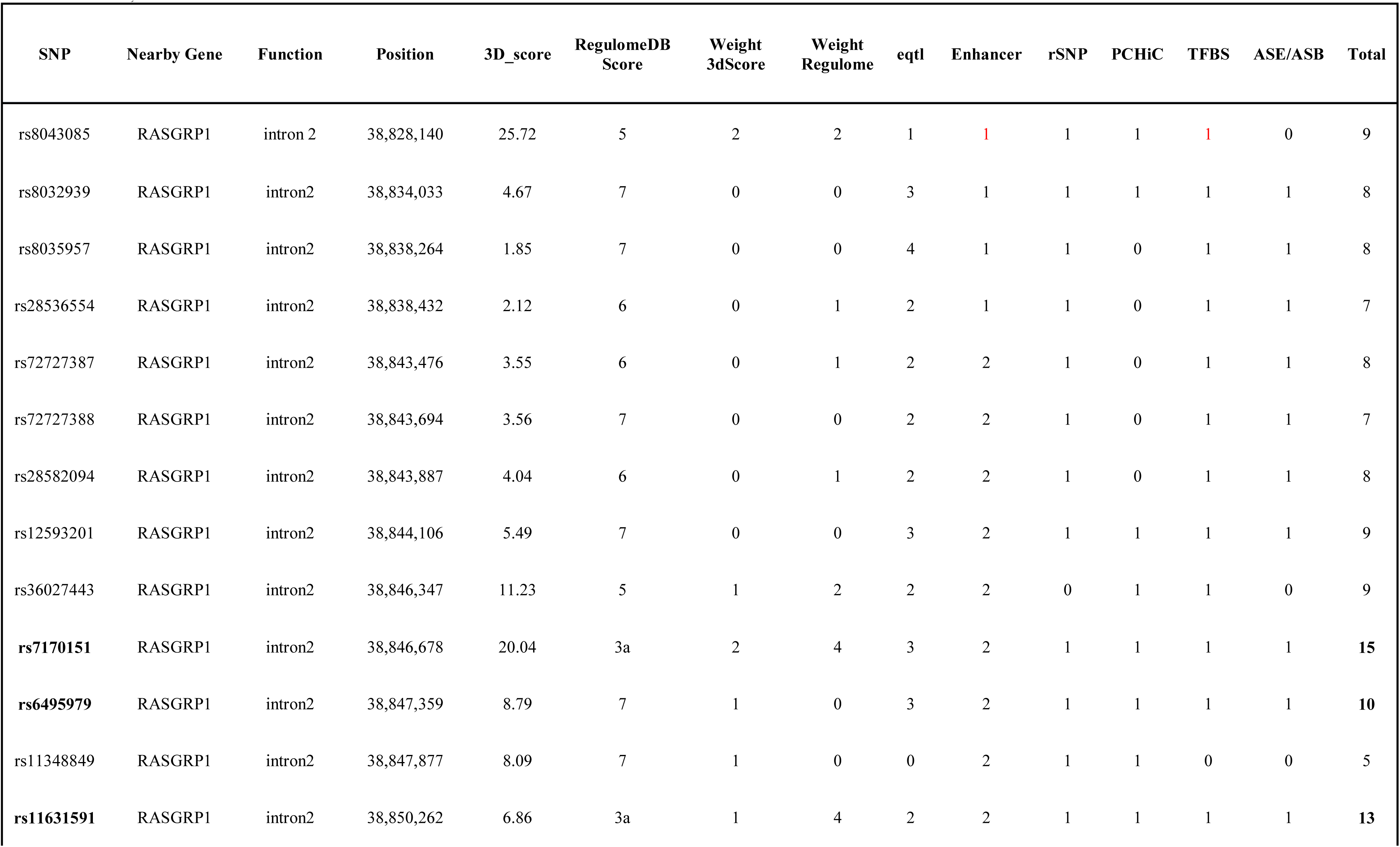
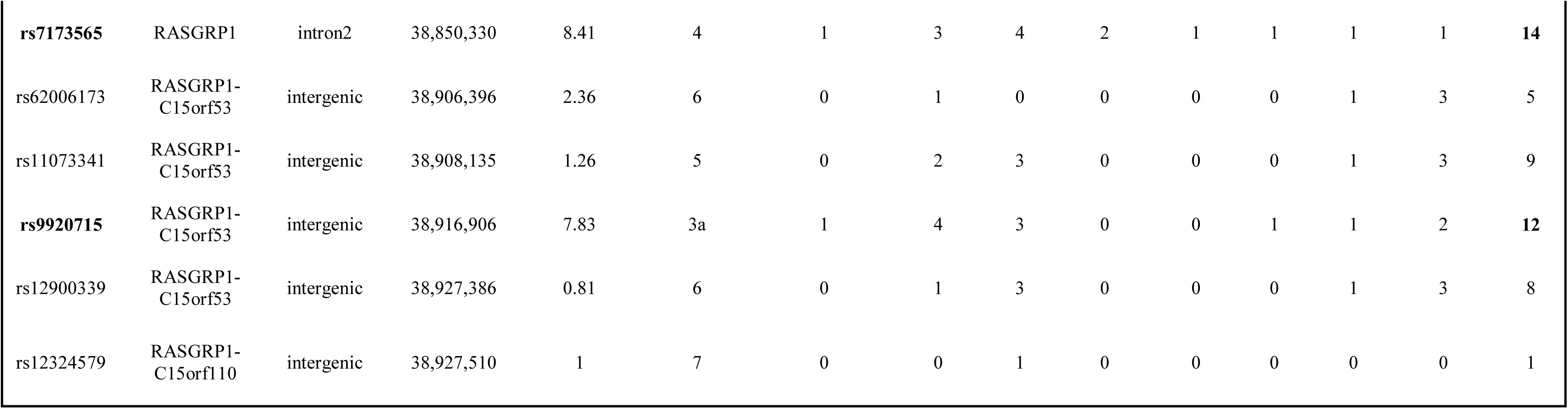
Relevant epigenetic features of genome-wide significant SNPs. We integrated scores from 3dSNP, RegulomeDB and rSNP with blood cell-specific information for eQTLs, enhancer/super-enhancer existence, promoter capture HI-C (PCHiC), transcription factor binding site (TFBS) disruption and allele-specific expression/binding (ASE/ASB) into a weighted score for SNP prioritization. We chose the top three SNPs for further validation (rs11631591 and rs7173565 were used together because of the short distance between them).

### Identification of DNA-binding proteins

#### Electrophoretic mobility shift assays (EMSAs) and DNA pulldown assays

Jurkat cell lines were obtained from ATCC and maintained in RPMI 1640 medium with 2 mm L-glutamine, 100 μg/ml each of streptomycin and penicillin, and 10% fetal bovine serum at 37°C with 5% CO_2_. Cells were harvested at a density of 8×10^5^ cells/ml, and nuclear extracts were prepared using the NER nuclear extraction kit (Invitrogen) with complete protease inhibitors (Roche Diagnostics). Protein concentrations were measured using a BCA reagent. Biotinylated DNA sequence surrounding the candidate SNPs (rs7170151 and rs11631591) was prepared using a synthetic single-stranded DNA sequence (Integrated DNA Technologies, USA) (**Supplementary Table 3**). Biotinylated DNA sequence with a 5-bp deletion at the SNP region served as a control for the assay. Twenty-five pmol of each DNA product was bound to 1 mg Dynabeads® M-280 Streptavidin (Invitrogen, USA) as per the manufacturer’s recommendations. Dynabeads M-280 Streptavidin (Dynal, Inc., Lake Success, NY, USA) were prepared by washing three times in phosphate-buffered saline (pH 7.4) containing 0.1% bovine serum albumin and two times with Tris-EDTA containing 1M NaCl. Between each wash, beads were pulled down with a Dynal magnetic particle concentrator. Double-stranded, biotinylated oligonucleotides were added to the washed beads, and the mix was rotated for 20–30 min at 21□°C. Equal cpm of proteins translated *in vitro* were diluted to 1x with binding buffer and mixed with ∼100 μg of Dynabeads containing 10 pmol of the individual oligonucleotide probe in a final volume of 250 μl. The mixture was rotated at room temperature for 20 min. Proteins bound to the beads were separated from unbound proteins by successive washes, three times with 0.5x binding buffer and once with 1x binding buffer. Higher stringency washes included two washes with 2x binding buffer. Beads and bound proteins were pulled down with a magnetic concentrator, suspended in 1x sample buffer, boiled for 5 min, and resolved on SDS-PAGE gels followed by peptide mass fingerprint MALDI-MS analysis of single bands.

#### Mass spectrometry analysis

Mass spectrometry analysis was performed using a Thermo-Scientific LTQ-XL mass spectrometer coupled to an Eksigent splitless nanoflow HPLC system. Bands of interest were excised from the silver nitrate-stained Bis-Tris gel and de-stained with Farmer’s reducer (50 mM sodium thiosulfate, 15 mM potassium ferricyanide). The proteins were reduced with dithiothreitol, alkylated with iodoacetamide, and digested with trypsin. Samples were injected onto a 10 cm × 75 mm inner diameter capillary column packed with Phenomenex Jupiter C18 reverse phase resin. The peptides were eluted into the mass spectrometer at a flow rate of 175 nl/min. The mass spectrometer was operated in a data-dependent mode acquiring one mass spectrum and four CID spectra per cycle. Data were analyzed by searching all acquired spectra against the human RefSeq databases using Mascot (Matrix Science Inc., Boston, MA, USA). Minimum identification criteria required two peptides with ion scores greater than 50% and were verified by manual inspection. We verified the identity of the assayed proteins by Western blot.

#### Confirmation of identified protein by Western blot

Mass spectrometry-identified proteins were confirmed by Western blot. Jurkat nuclear extracts after DNA pulldown assay were lysed in sample buffer [62.5 mM Tris·HCl (pH 6.8 at 25°C), 2% wt/vol SDS, 10% glycerol, 50 mM dithiothreitol, 0.01% wt/vol bromophenol blue]. Equal amounts of protein were loaded onto a 10% SDS-PAGE gel (GTX gel BioRad USA). After it resolved, samples were blotted to Nitrocellulose paper using the Trans-blot Turbo Transfer System (BioRad, USA). Membranes were blocked using LI-COR blocking buffer for 2 hours and then incubated with primary antibody 1:1000 dilution (hnRNP-K, Santa Cruz USA) at 4°C overnight, and with a donkey anti-mouse IR-Dye 800 (LI-COR, USA) secondary antibody for 1 h. The membrane was imaged with a LI-COR Odyssey using Auto-Scan. Background-subtracted signal intensity was quantified using Image Studio 4.0 software.

#### Chromatin immuno-precipitation (ChIP) assay followed by qPCR (ChiP-qPCR)

ChIP assays were performed using the MAGnify ChIP system (Life Technologies, NY) according to the manufacturer’s protocol. Jurkat cells were fixed for 10 min with 1% formaldehyde to crosslink DNA-protein and protein-protein complexes. The cross-linking reaction was stopped using 1.25 M glycine for 5 min. The cells were lysed, sonicated to shear DNA and sedimented. Then, their diluted supernatants were incubated with 5 μg hnRNP-K antibody. Ten percent of the diluted supernatants were saved as “input” for normalization. Several washing steps were followed by protein digestion using proteinase K. Reverse crosslinking was carried out at 65°C. DNA was subsequently purified and amplified by quantitative PCR on an SDS 7900 (Applied Biosystems) using specific primers. Because the Jurkat cell line is heterozygous for the SNPs rs11631591 and rs7170151, we performed Sanger DNA sequencing with the ChIP-eluted PCR product.

#### Isolation of CD3^+^ T-cells from human blood

We used leukoreduction system chambers (LRS chambers) from human blood donors. LRS chambers were obtained from the Oklahoma Blood Institute (OK, USA) (**Supplementary Table 12**, **Supplementary Figure 9**). LRSCs were sterilized externally using 70% (v/v) ethanol and handled in a class 2 laminar flow cabinet. External tubing was cut, the chamber inverted over a 50 ml sterile centrifuge tube (Greiner Bio-One) and the contents allowed to drip through. The contents (usually 20 ml) were then diluted to 90 ml in RPMI medium. The peripheral blood mononuclear cells (PBMCs) were isolated by carefully layering 30 ml fractions over 17 ml of histopaque-1077 (Sigma-Aldrich), which was then centrifuged at 340 g for 45 min at 20°C. The PBMC layer was isolated and washed three times with culture medium with cells centrifuged at 340 g for 15 min for the first wash and 10 min for the subsequent two washes. The isolated PBMCs were counted and viability assessed with Trypan blue using a hemocytometer, then centrifuged at 340 g for 10 min. The untouched CD3+ T cells were collected using MojoSort™ Human CD3^+^ T-Cell Isolation Kit as per manufacture instructions (BioLegend, San Diego, CA).

#### Inhibition of hnRNP-K and ERK phosphorylation

Inhibition of hnRNP-K was performed in CD3^+^ T cells from healthy controls, as well as in Jurkat T-cells using 5-Fluorouracil (5-FU) (Sigma Aldrich, USA) as described previously [54]. Isolated CD3^+^ T-cells and Jurkat cells were cultured in RPMI-1640 medium containing 10% heat-inactivated fetal bovine serum (Invitrogen) and kept at 37°C in 5% CO_2_ conditions. For 5-FU treatment, the drug was first dissolved in dimethyl sulfoxide (DMSO) and further diluted in medium before use. Cells were treated with 20ng/μl 5-FU unless otherwise stated. Next, to examine whether hnRNP-K and/or *RASGRP1* down-regulation by 5-FU led to inhibition of EKR phosphorylation of ERK, Jurkat and CD3^+^ T-cells were pretreated with PMA 5ug/μl for 30 minutes, prior to drug (5-FU) treatment. Inhibition of hnRNP-K and *RASGRP1* was detected using mRNA expression analysis with quantitative PCR (after 48 hours) and by Western blot (after 72 hours).

## Results

### Patients and samples

We used five Asian cohorts and one cohort of European descent; sample sizes for the meta-analysis were 9,529 SLE cases and 22,462 controls (**Table 1**).

### Fine-mapping, replication and meta-analysis of RASGRP1 association

First, we probed our previously reported SLE-associated region (chr15: 38.4-39.2 MB, hg19) and extracted association results for six cohorts from the region containing the genes *RASGRP1* (RAS guanyl-releasing protein 1, a diacylglycerol-regulated guanine nucleotide exchange factor) and *C15orf53* (encoding a protein of unknown function linked to alcohol dependence [55]). The strongest association signal among Asian cohorts localized to intron 2 of *RASGRP1* **(Figure 2**, **Table 2**). Meta-analysis with all cohorts identified the largest signal at intronic SNP rs8032939 (P_meta_=3.2×10^−11^, OR (95%CI) = 0.88 (0.85-0.92)). We identified 17 genome-wide significant (GWS) SNPs (P_meta_<5×10^−8^). Our previously reported lead SNP rs12900339 [4] did not reach GWS (P_meta_=9.2×10^−7^) (**Table 2**). Analysis of the association signals in the context of linkage disequilibrium (LD) of 1000Genome populations (EUR, ASN; **Figure 2**) identified two uncorrelated association signals (**Supplementary Table 1**). The main signal occurred at rs8032939 in intron 2 (**Figure 2**), while the second signal localized to the intergenic region between *RASGRP1* and *C15orf53*: SNP rs9920715 (60 kb 5’ of *RASGRP1*; P_meta_ = 5.1×10^−9^; OR (95%CI) = 0.89(0.86-0.93)). Many (27 of 119 SNPs) variants were intronic (**Figure 2**). We then examined the 18 GWS SNPs with bioinformatic and epigenomic analysis (**Table 2**). Our top SNP (rs8032939) was previously reported as a rheumatoid arthritis (RA)-associated SNP [56]. Within the intronic signal, we also identified rs8035957 (P_meta_=1.3×10^−10^), associated with Type I Diabetes [57].

**Figure 2.**
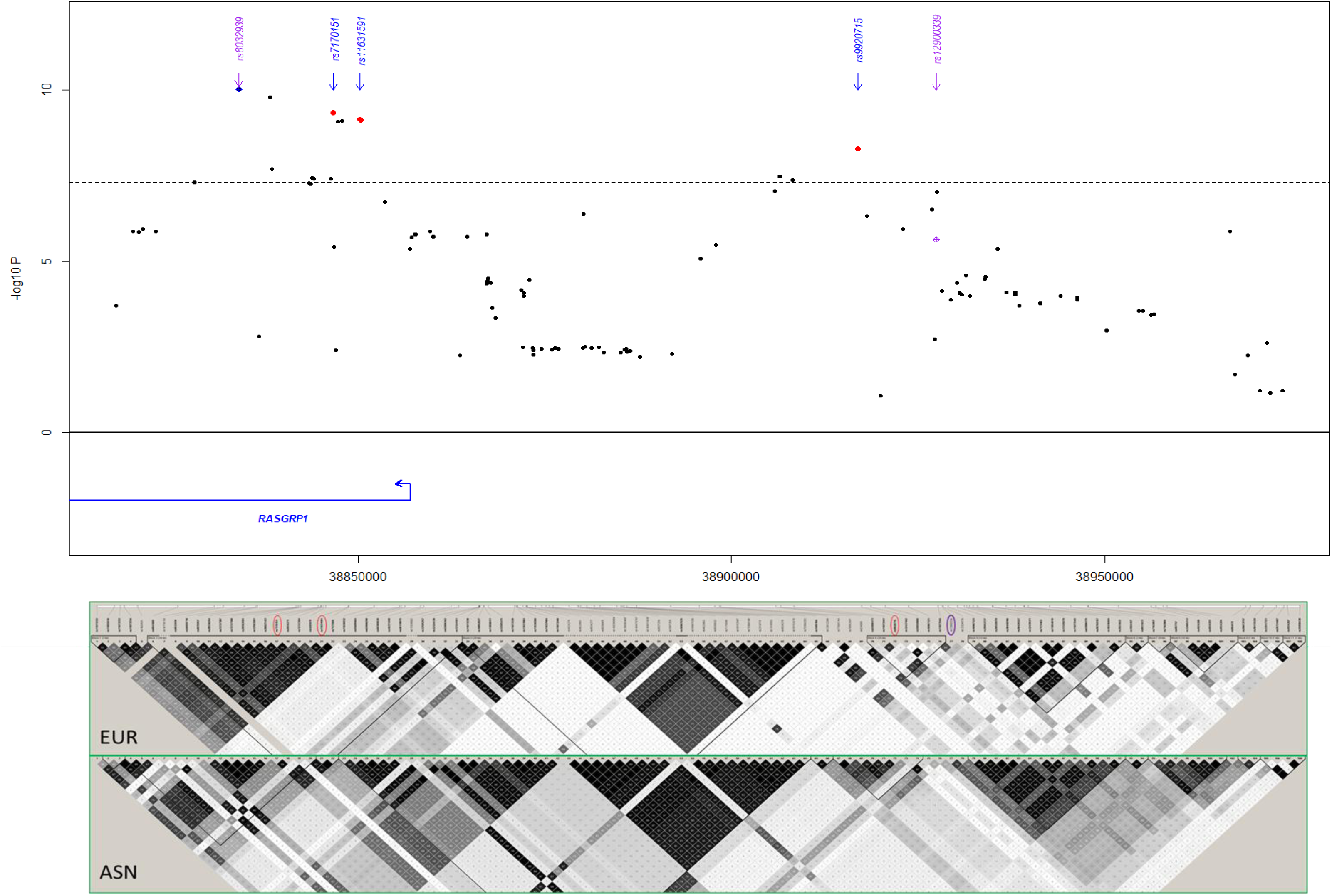
Meta-analysis in the *RASGRP1* region. Blue diamond: lead SNP rs8032939 following initial meta-analysis. Red circles: SNPs chosen for experimental validation. rs11631591-rs7173565 are considered together due to their proximity; only rs11631591 is labeled. Purple diamond: our previously reported[4] lead SNP rs12900339. Linkage disequilibrium in the region (bottom) is notably different between European (EUR) and Asian (ASN) populations.

### Evaluating functional SNPs

To identify putative functional SLE SNPs in and around *RASGRP1*, we computed weighted scores for each SNP by integrating multiple sources of functional annotation, including allele-dependent gene expression, overlap with annotated enhancers and promoters, binding affinity to transcription factors and collocation with anchors in promoter-enhancer-capture Hi-C experiments (**Supplementary Table 2**).

### Gene expression

We then identified allele-dependent changes in gene expression by annotating SNPs using expression quantitative trait locus (eQTL) databases for multiple tissues (**Methods**). All candidate LD SNPs were eQTLs in blood cell lines (3.2×10^−3^>P>1.9×10^−4^; **Supplementary Table 4**), as well as in skin, esophagus, and testis (**Table 3**). The intronic (main signal) SNPs affected expression of both *RASGRP1* and *C15orf53*, while the intergenic (secondary) SNPs (in LD with rs9920715) altered expression of only *RASGRP1*. *RASGRP1* SNPs also affected expression of long non-coding RNAs (lncRNAs) *RP11-102L12.2* and *RP11-275I4.2* in non-blood cell lines. All eQTL risk alleles increased expression of *RASGRP1* in multiple cell lines (**Supplementary Table 4**, **Supplementary Figure 2**), but had opposing effects on the neighboring gene *C15orf53* (**Supplementary Table 4)**. We also found significant effects of two linked SNPs (rs11073344, rs11631591) on methylation of *RASGRP1* in T-cells and neutrophils, respectively (**Supplementary Table 5**).

### Overlap with enhancers and super-enhancers

Then, we investigated the potential of the candidate SNPs to act as enhancers of *RASGRP1* expression. Three GWS SNPs (rs6495979, rs11631591, and rs7173565) overlapped with ENCODE-annotated enhancers for *RASGRP1* in lymphoblastoid cells (GM12878, GM12892) and also in CD8^+^ T-cells. These three GWS SNPs (all intronic) localized to super-enhancers (*i.e.* collections of multiple contiguous enhancers [58]) for *RASGRP1* in CD4^+^ CD25^−^ CD45RA^+^ naïve cells, CD4^+^ CD25^−^ CD45RO^+^ memory cells, CD8^+^ primary cells, CD4^+^ CD25^−^ Il17^+^ phorbol myristate acetate (PMA)-stimulated Th17 cells, and CD4^+^ CD25^−^ Il17^−^ PMA-stimulated Th17 cells; **Supplementary Table 6**). This suggests that these SNPs may regulate *RASGRP1* in T-lymphocytes.

### Chromatin interactions

Since all candidate SNPs reside outside of the *RASGRP1* promoter, we investigated if the SNPs overlapped with anchors in promoter-enhancer connections though chromatin interactions. We used promoter-capture Hi-C data on blood cell lines, in particular T-cells, to identify physical interactions between the intronic signal and the *RASGRP1* promoter (**Supplementary Figure 3**, **Supplementary Table 7**). We also examined the physical interaction between the intergenic region (represented by rs9920715) and the promoters of *RASGRP1* and *C15orf53*. We identified multiple significant promoter-enhancer interactions between the intronic signal and *RASGRP1, C15orf53, FAM98B* and *SPRED1*, and multiple interactions between the intergenic signal and the promoter of *RASGRP1* (**Supplementary Table 7**).

**Figure 3.**
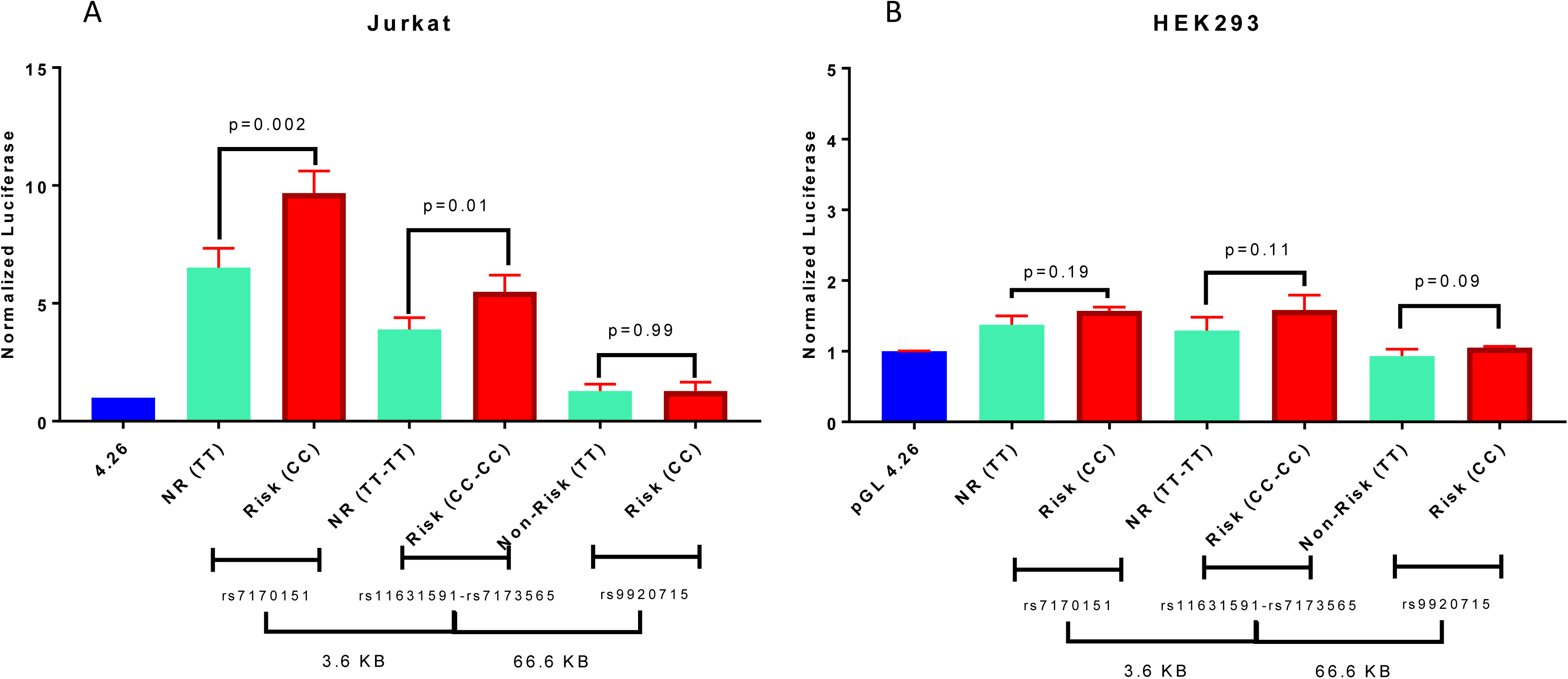
Luciferase reporter assay for rs7170151, rs11631591-rs7173565 and rs9920715. A. Jurkat cells. B. HEK293 cells. Empty vector pGL4.26 was used as reference. NR: non-risk. P-values are for Student’s t-test.

### Effect on cytokine production

A critical feature in SLE pathogenicity is cytokine production [59]; thus, we investigated if these SNPs alter cytokine abundance. Our candidate SNPs significantly increased expression of interleukins IL6 and IL22 and tumor necrosis factor (TNFα), while SNP rs9920715 exclusively increased IL22 expression (**Supplementary Table 8**).

### Allele-specific binding

We found that 14 of the candidate GWS SNPs also had allele-specific binding (ASB) to H2K27ac in monocytes, neutrophils, and T-cells (**Supplementary Table 9**), while rs9920715 showed ASB with H3K4me1 in T-cells and neutrophils. To characterize the regulatory mechanisms involved, we assessed ASB of histone marks H3K4me1 and H3K4me3 at and around candidate SNPs (**Supplementary Table 9**, **Table 3**). We identified a significant regulatory region associated with promoter mark H3K4me3 with a higher binding affinity to the extended region (∼1 kb) containing the risk alleles (both C) of intronic SNPs rs11631591-rs7173565 (**Supplementary Figure 4a**). In addition, we identified marginally significant ASB to enhancer mark H3K4me1 at SNPs rs6495979 and rs7170151, which tagged a regulatory region within ∼500 bp (**Supplementary Figure 4b**). These data indicate that allele-specific differences might affect chromatin interactions.

**Figure 4.**
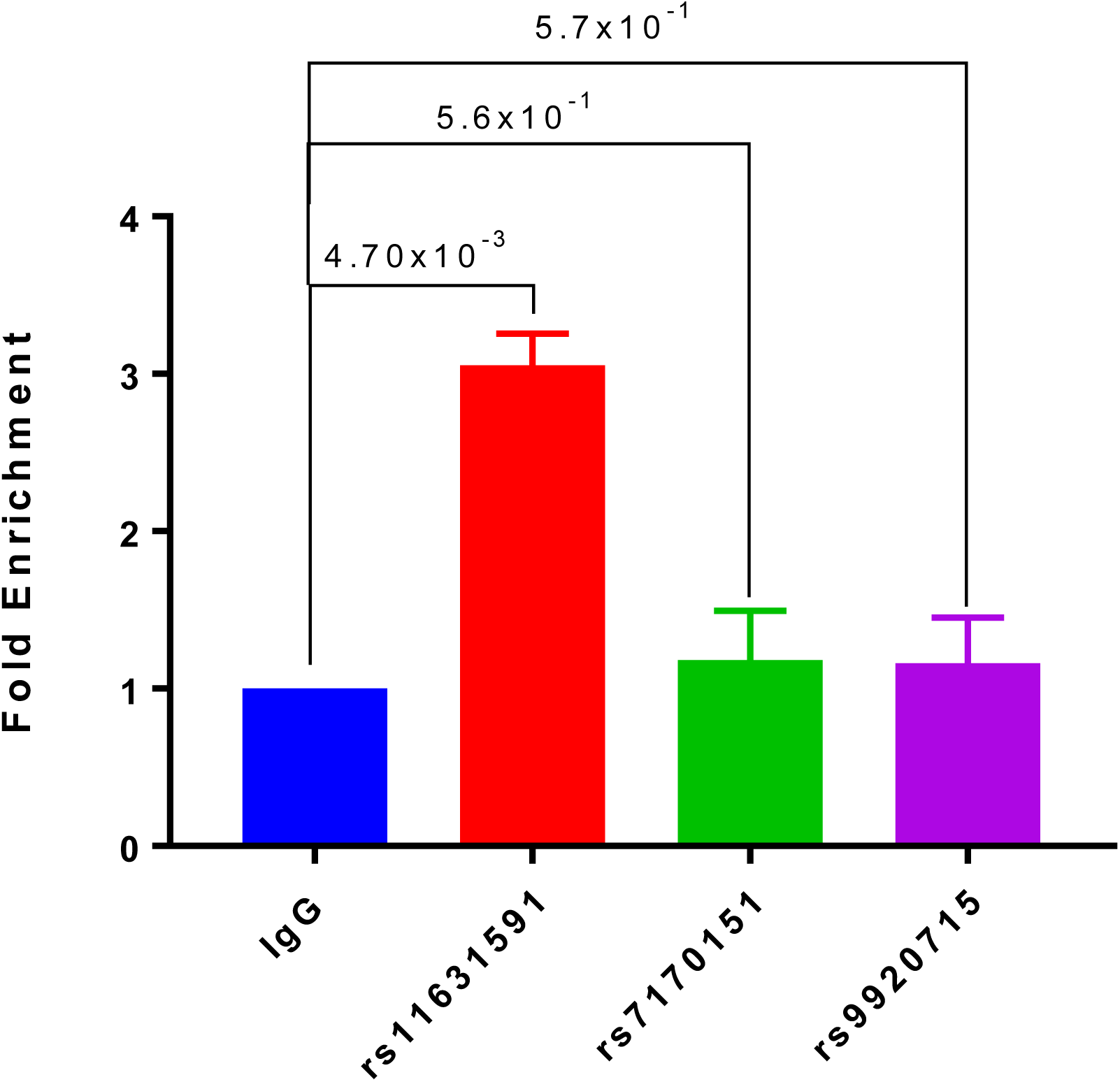
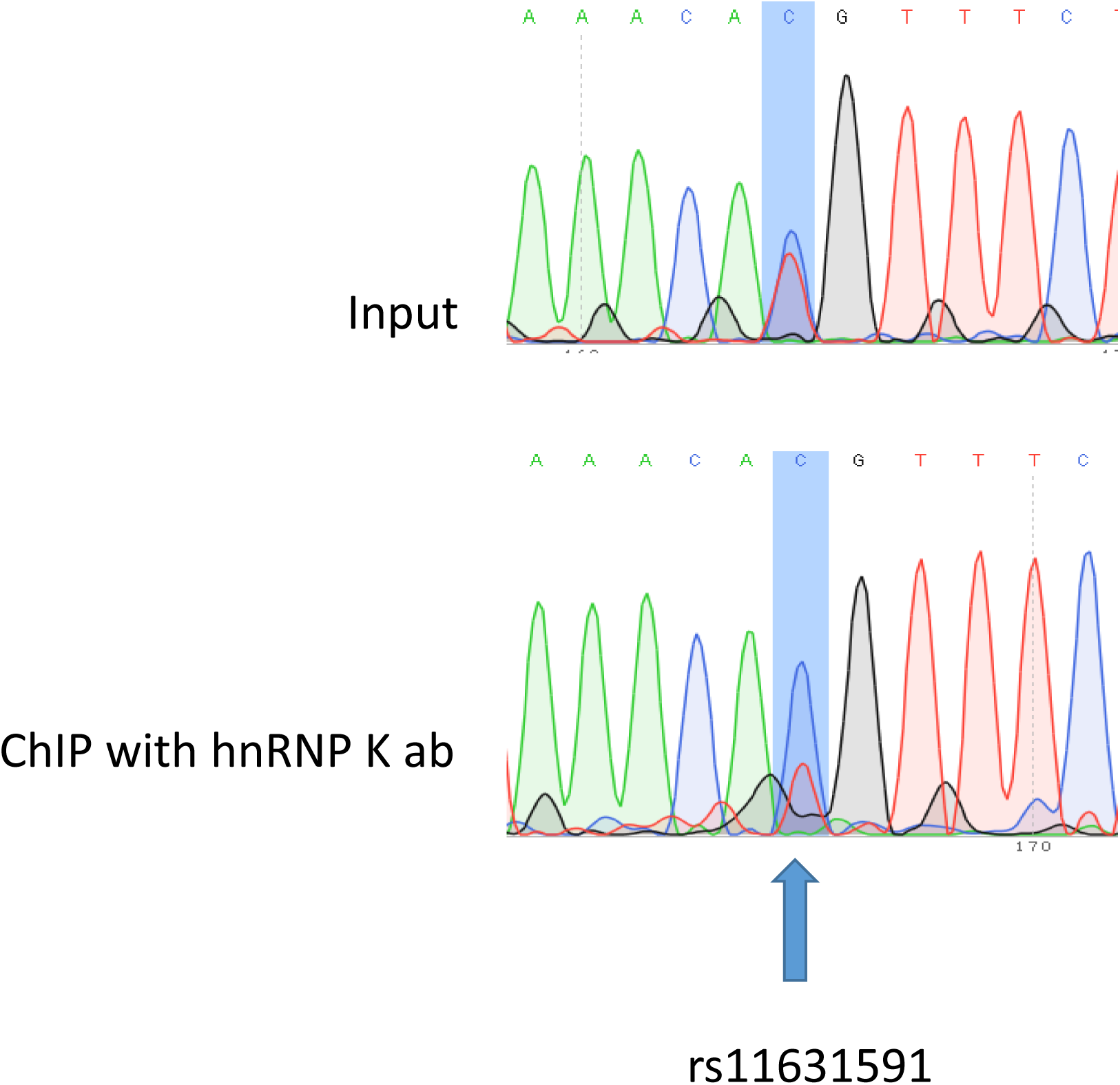
**A**. ChIP-qPCR of sequences containing SNPs rs11631591-rs7173565, rs7170151 or rs9920715 in Jurkat cells. SNP rs11631591 showed 3-fold enrichment of hnRNP-K binding over IgG control. No significant enrichment at the other two SNPs was observed. P-values are for Student’s t-test. **B**. Sequence chromatographs showing the difference in zygosity of rs11631591 between the input (equal binding to the two alleles, above) and the ChIP assay at the risk allele (2-3x more binding to the risk C allele, below).

### Validation of enhancer by luciferase assays

When testing in a luciferase reporter assay, rs7170151 and rs11631591 showed marked (up to 10-fold over empty vector) enhancer activity in Jurkat cells (P=3.0×10^−4^, P=1.0×10^−3^, respectively) and less so (1.6-fold) in HEK293 cells (P=4.0×10^−2^, P=3.0×10^−3^); however, rs9920715 functioned as a very weak enhancer only in HEK293 (P=4.1×10^−2^) (**Figure 3**). Furthermore, rs7170151 and rs11631591 showed dramatic allelic differences in enhancer function. Genomic regions containing homozygous risk alleles of rs7170151 (C) and rs11631591 (C) showed significantly higher enhancer activity (∼50% increase; P=1.0×10^−2^ and P=2.3×10^−3^, respectively; **Figure 3A**) compared to non-risk alleles, but only in Jurkat cells. This allele-dependent enhancer activity is consistent with the allele-specific expression we observed in the eQTL data. There were no significant differences in HEK293 cell lines (**Figure 3B**), suggesting that enhancer activity depends on white blood cell-specific factors. The third intergenic SNP (rs9920715) did not show enhancer activity in any assayed cell type (**Figure 3A**, **B**).

### Transcription factor binding

We next assessed allele-specific changes in transcription factor binding site (TFBS) affinity using five motif databases (**Methods**). We identified 256 TFBSs significantly affected by ten of our SNPs (**Supplementary Table 10)**. Notably, we found 43-fold higher affinity of promoter-specific TF YY1 to the non-risk allele (T) of rs7173565 and 42-fold higher affinity of TF GATA (GATA1..3.p2 motif) to the risk (T) allele of rs6495979. Interestingly, SLE-risk ETS1[60] binding had 10-fold higher affinity to the risk (C) allele of rs7173565, while SLE-risk IRF5[61] bound 6-fold more tightly to the non-risk (C) allele of rs6495979.

### Identification of DNA-binding proteins

We detected DNA-binding protein complexes using electrophoretic mobility shift assays (EMSAs) and DNA pulldown assays using a 41 bp-long dsDNA containing the rs11631591-rs7173565 (homozygous risk, CC; or homozygous non-risk, TT) alleles (**Supplementary Table 11**). We prepared nuclear extracts from Jurkat cells and incubated them with biotin-labeled dsDNA (risk *vs*. non-risk) bound to magnetic beads coated in streptavidin. EMSA showed multiple bands of DNA-bound proteins (**Supplementary Figure 5**). We observed allele-specific binding of a protein complex at 75 kDa. Although EMSA is not a quantitative assay, we observed in multiple independent experiments that the intensity of the band with the risk (CC) oligo was darker than with the non-risk (TT) oligo, suggesting allele-specific differential binding (**Supplementary Figure 5**). Using mass spectrometry analysis of bound proteins, we identified heterogeneous nuclear ribonucleoprotein K (hnRNP-K) isoform b as the most abundant bound protein (**Supplementary Table 11**). hnRNP-K was also the protein whose binding was most diminished by substitution of the risk CC by non-risk TT nucleotides. We also confirmed that the identified protein bound with the risk oligo for the region of rs11631591 was hnRNP-K through EMSA followed by Western blot (**Supplementary Figure 6**).

**Figure 5.**
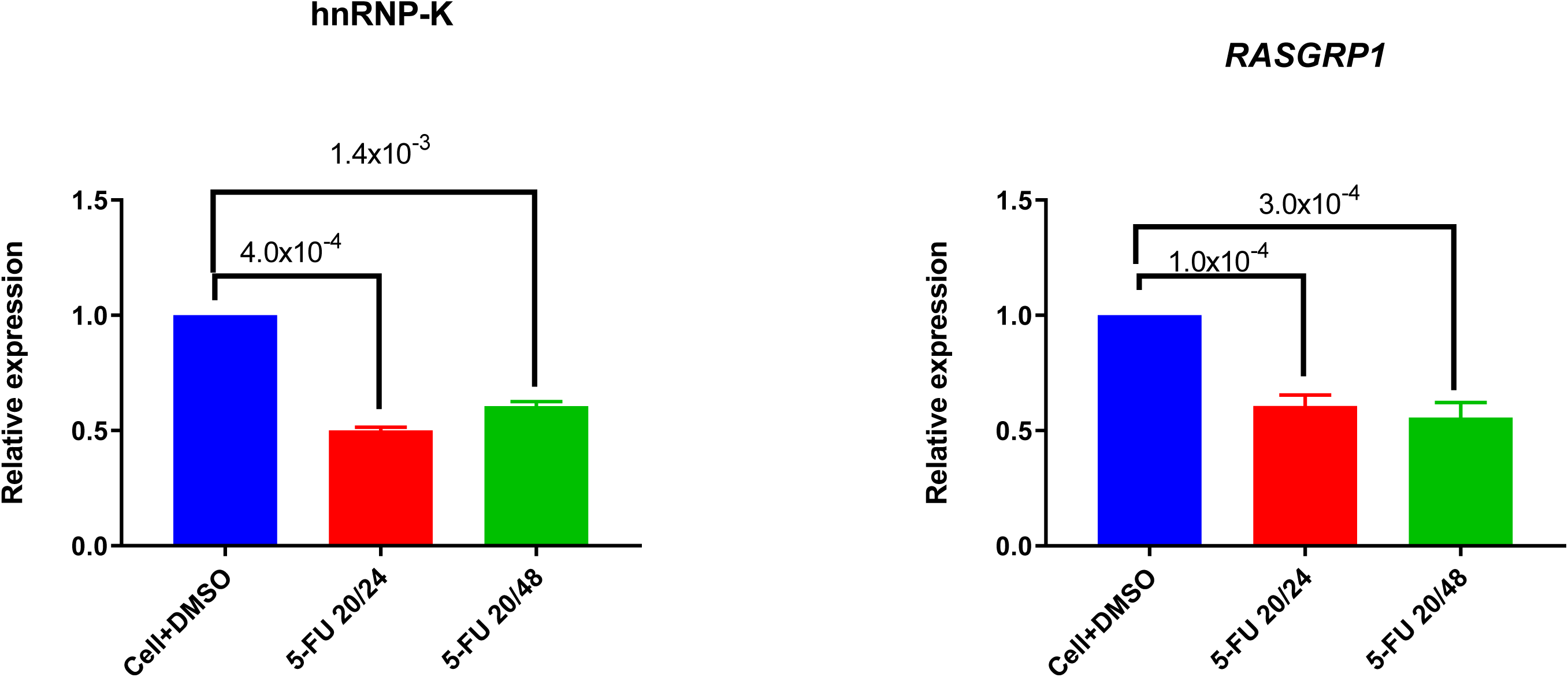
Downregulation of hnRNP-K by 5-FU treatment. 5-FU treatment reduces hnRNP-K expression levels in Jurkat cells. Jurkat cells were treated with DMSO vehicle or 5-FU (20 ng/μl) for 24 or 48 hours. *hnRNP-K* (A) and *RASGRP1* (B) were examined with *GADPH* as loading control.

**Figure 6.**
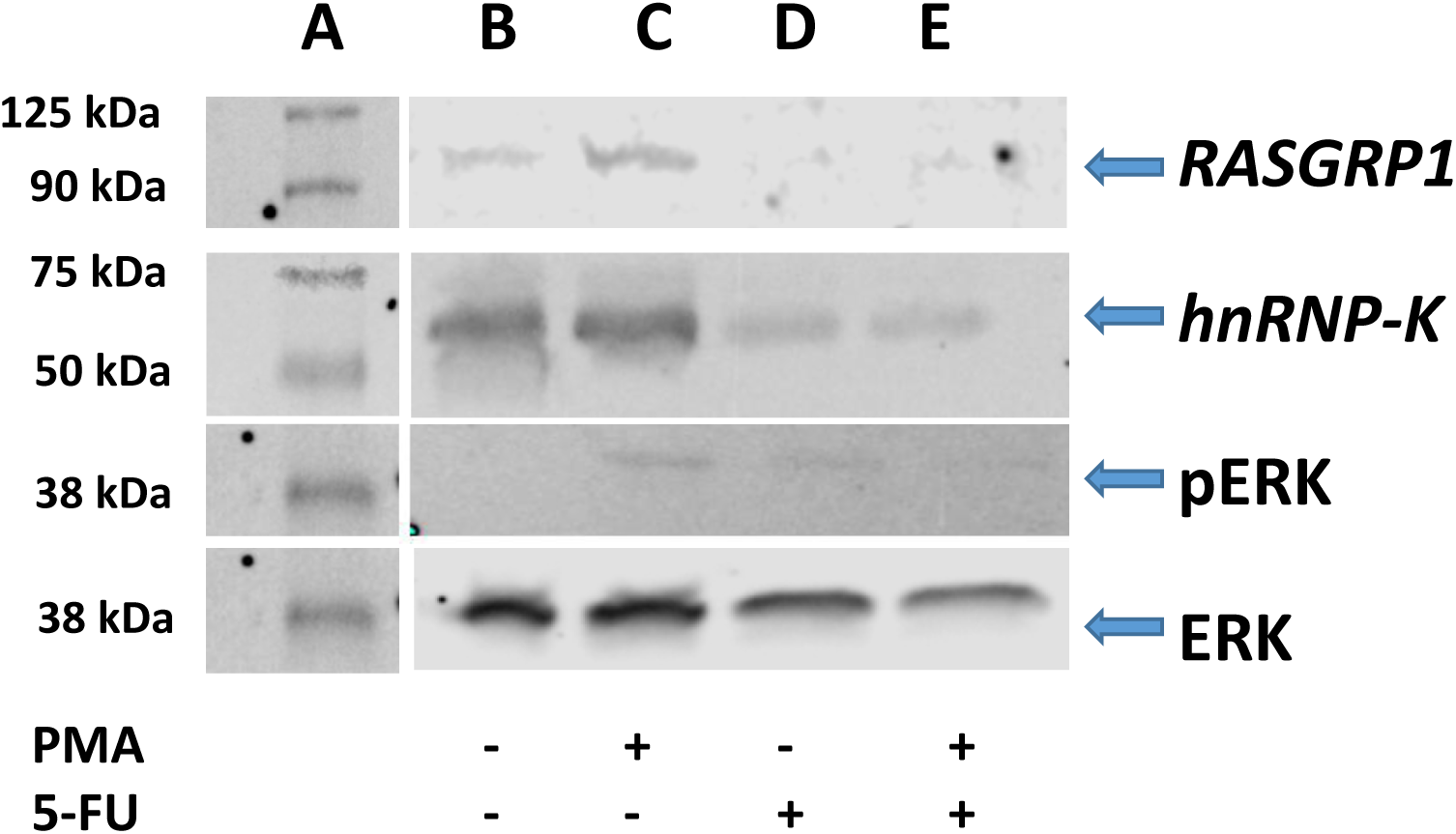
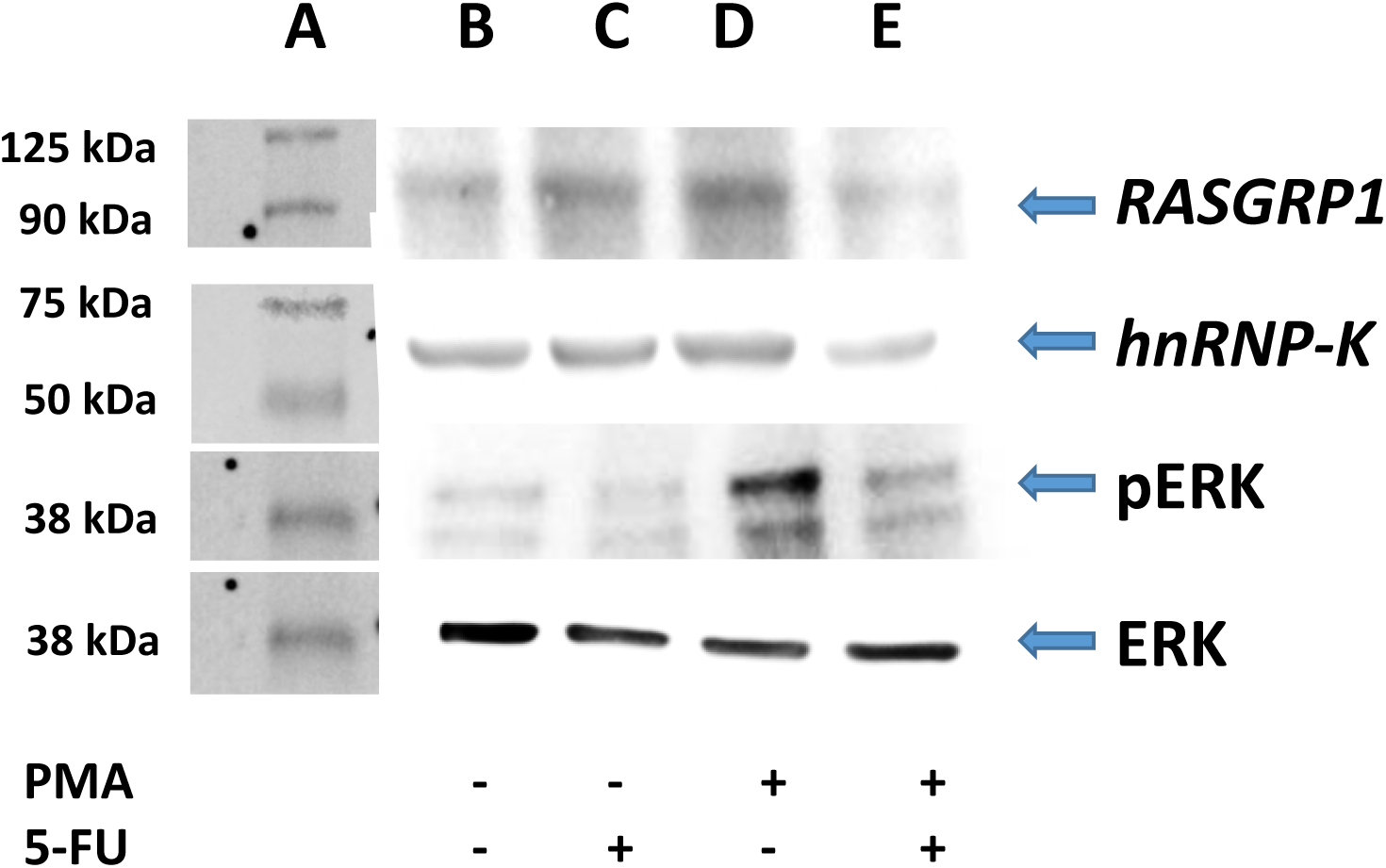
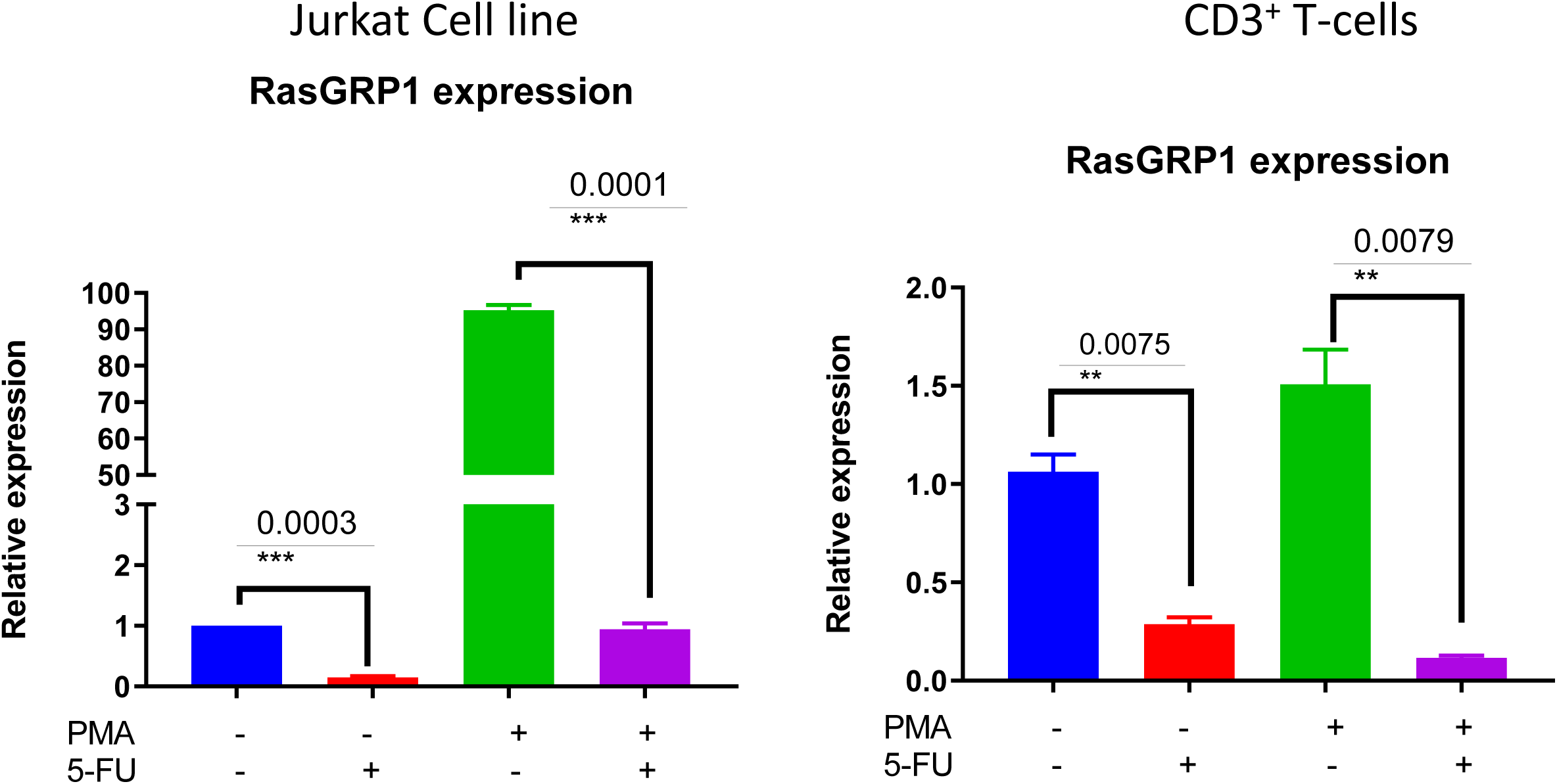
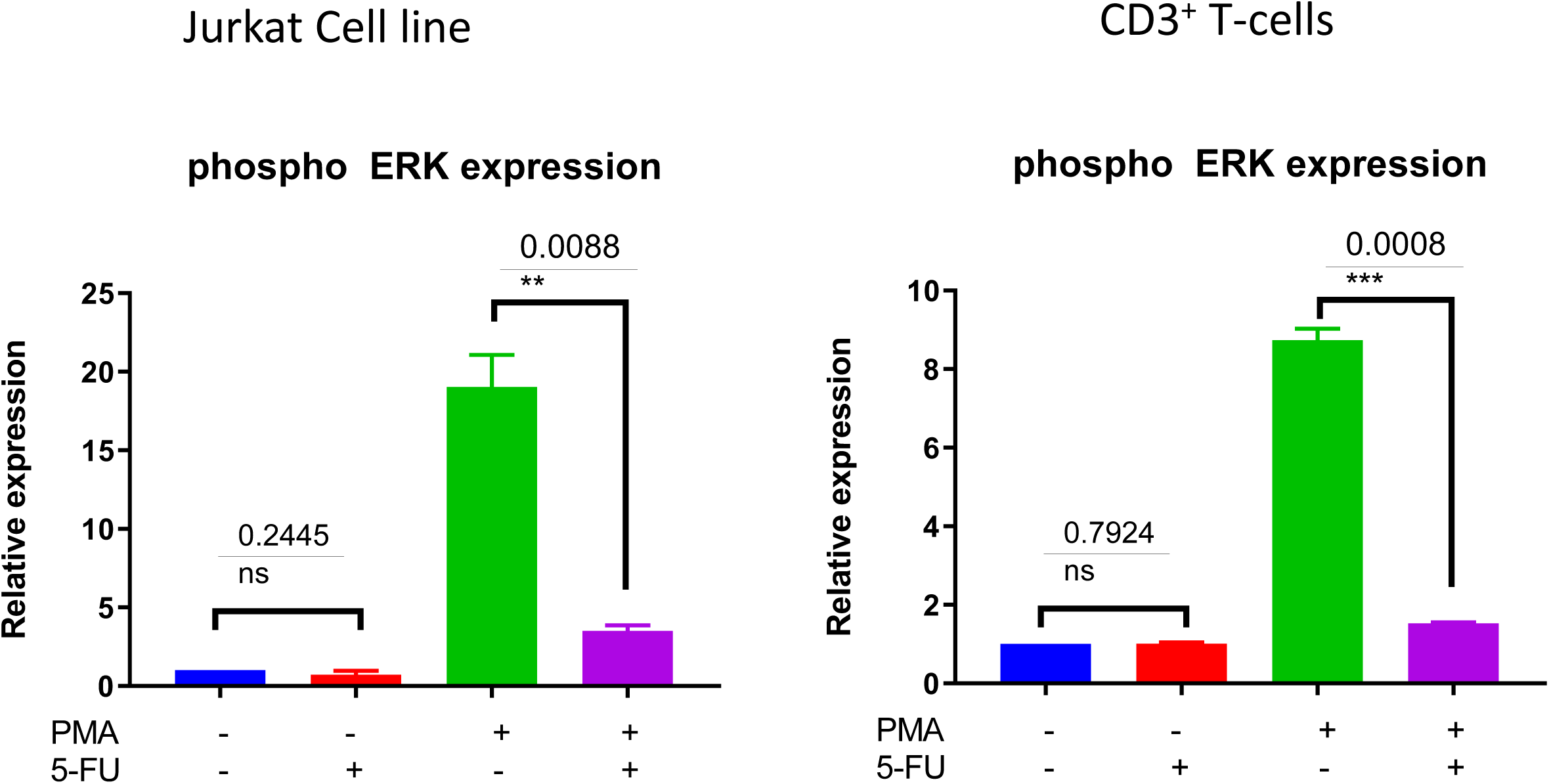
**A**. *RASGRP1* reduction influences the phosphorylation of ERK. 5-FU treatment reduces hnRNP-K and *RASGRP1* expression levels in Jurkat and healthy human CD3^+^ T cells. Pretreatment with PMA increases levels of *RASGRP1* and phospho-ERK. Inhibition of hnRNP-K with 5-FU decreases levels of *RASGRP1* and phospho-ERK, even after PMA stimulation. **B**. 5-FU treatment reduces hnRNP-K as well as *RASGRP1* expression level in primary CD3^+^ T-cells. Pretreatment with PMA induces *RASGRP1* expression and leads to phosphorylation of ERK and reduction of *RASGRP1*; treatment with 5-FU also leads to reduction of phosphorylation of ERK. **C**. Densitometric analysis for RasGRP1 normalized to β-actin: primary T-cells and Jurkat cells. Results are presented as relative fold-change following drug treatment with and without stimulation. **D**. Densitometry analysis for phospho-ERK normalized to β-actin: primary T-cells and Jurkat cells. Results are presented as relative fold-change following drug treatment with and without PMA stimulation.

### SNPs bind to different transcription factors in an allele-specific manner

Using EMSA and mass spectrometry, we showed that hnRNP-K protein has tighter binding affinity to the risk genotype (CC) of SNP rs11631591-rs7173565. We validated these findings using Jurkat (heterozygous CT at rs11631591-rs7173565) to perform chromatin-immunoprecipitation (ChIP) followed by RT-qPCR (ChIP-qPCR). We observed significant enrichment in binding of the hnRNP-K antibody to the SNP region of rs11631591, but did not observe any binding of hnRNP-K antibody to either rs7170151 or rs9920715 (**Figure 4A**). To determine preferential or allele-specific binding, we performed Sanger sequencing on the region containing rs11631591-rs7173565. Both alleles were present in the original input sample; however, only the risk allele (C) was detected significantly higher than the non-risk allele (T) in chromatograms of the ChIP-eluted PCR product (**Figure 4B**). These data suggest preferential allele-specific binding of the rs11631591-rs7173565 risk locus to hnRNP-K.

### hnRNP-K plays an important role in *RASGRP1* expression

To investigate the role of endogenous hnRNP-K in Jurkat and primary CD3^+^ T-cells, we transiently inhibited hnRNP-K using 5-fluorouracil (5-FU). After 5-FU treatment (48 hours), we observed significantly reduced mRNA expression for both hnRNP-K (P=1.4×10^−3^; **Figure 5A**) and *RASGRP1* (P=3.0×10^−4^; **Figure 5B**). 5-FU-induced hnRNP-K downregulation correlated with reduced expression of *RASGRP1* (**Figure 6A**, **B**). This result suggests that hnRNP-K plays an important role in *RASGRP1* expression in Jurkat cells as well as in primary T-cells. Furthermore, we observed the reduction of ERK phosphorylation with 5-FU after initial induction with PMA in Jurkat and primary T-cells (**Figure 6 A-D**). It is of note that stimulation with PMA did not influence cell viability (**Supplementary Figure 7**).

## Discussion

In this study, we fine-mapped our previously reported SLE locus near *RAS guanyl-releasing protein 1* (*RASGRP1*), a lynchpin of T-cell development and the RAS/MAP kinase signaling cascade following antigen exposure. We performed a trans-ethnic meta-analysis of the locus with cohorts of Asian and European descent, followed by multiple lines of bioinformatic analysis of its epigenetic context to prioritize SNPs as candidate causal variants. Experimental testing of the top candidates validated them as plausible variants underlying association of this locus with SLE (and perhaps other autoimmune phenotypes).

We identified two independently associated regions correlated with *RASGRP1* regulation and expression. The first signal lies in *RASGRP1* intron 2, represented by SNPs rs11631591-rs7173565 and rs7170151, which regulate *RASGRP1* expression as eQTLs (esophageal mucosa and skin), enhancers (in CD8^+^ T-cells, and thymic and lymphoblastoid cell lines), and as interaction anchors with the nearby *C15orf53* promoter. The SNPs in this region are within a robust enhancer, with the risk alleles (rs7170151-C and rs11631591-C/rs7173565-C) greatly increasing *RASGRP1* expression in multiple tissues (databases) and in Jurkat T-cells (our experiments). Furthermore, this enhancer is targeted by promoter interactions in CD8^+^ and CD4^+^ T-cells, B-cells, and monocytes [62] (**Figure 3**). We also identified another intergenic signal around 60 kb 5’ of *RASGRP1*, at rs9920715, another SNP within promoter-interacting chromatin that acts as an eQTL for *RASGRP1* in B- and T-cell lines [62]. However, this SNP did not show enhancer activity in our assays.

Mammalian gene regulatory elements, especially those that are tissue-specific, show high *in vivo* nucleosome occupancy, which can effectively compete with TF binding [63, 64]. This nucleosome-mediated restricted access to regulatory information is a key element for inducible or cell type-specific control of gene expression [65]. In the current study, we observed strong enhancer activity at rs11631591-rs7173565 or rs7170151 only in Jurkat but not HEK293 cells. Furthermore, our candidate SNPs show allele-specific *RASGRP1* expression, with the risk alleles driving substantially more (∼50%) expression than the non-risk alleles. Other studies on numerous complex diseases have demonstrated enrichment of disease-associated loci in cell type-specific regulatory regions of corresponding disease-relevant cell types [58, 66-69]. Additional studies now document the direct effects of common variation in enhancer elements on enhancer states [70-73], gene expression [70, 74], and disease [75-79]. Risk alleles of rs11631591 also showed significant binding to hnRNP-K protein in an allele-specific manner.

DNA/protein interaction assays demonstrated that hnRNP-K preferentially binds to sequences containing the rs11631591 risk (C) allele. We confirmed this allele-specific binding by EMSA and ChIP DNA sequencing. We only observed allele-specific binding of hnRNP-K at SNP rs116311591-rs7173565, but not at rs7170151 or rs9920715. We also observed that inhibition of hnRNP-K correlates with *RASGRP1* expression and ERK phosphorylation. In fact, expression of *RASGRP1* and *hnRNP-K* (P=9.8×10^−5^; P=1.4×10^−2^, respectively) in spleen (**Supplementary Figure 8**), shows a positive correlation between the risk allele of rs116311591 and both these genes. These data suggest that SNP rs11631591 is a functional SNP and may directly contribute to modulating *RASGRP1* expression. Abnormal expression of *RASGRP1* isoforms will perturb lymphocytes of SLE patients regardless of their clinical disease activity, and may contribute to impaired lymphocyte function and increased apoptosis in SLE patients [19]. Abnormal *RASGRP1* expression also induces ERK and JNK phosphorylation in the MAPK pathway, which in turn alters T-cell development, contributes to long-term organ damage and ultimately increases SLE susceptibility [22, 24, 25]. In the present study we also observed the role of *RASGRP1* expression in the phosphorylation of ERK activity. Altogether, our results indicate increased *RASGRP1* expression correlates with the risk alleles in our functional SLE loci and T-cell dysfunction. However, our study did not examine the differences in *RASGRP1* isoform expression reportedly associated with SLE and correlated with low *RASGRP1* expression [19].

In this study, we characterized the genetic risk of SLE in *RASGRP1*. We also propose a mechanism by which functional SNPs could affect SLE pathogenesis. We identified two functional regions affecting expression and regulation of *RASGRP1* in an intronic region including two SNPs (rs11631591 and rs7170151) and another in an intergenic region harboring SNP rs9920715. All identified SNPs are *RASGRP1* eQTLs and exhibit regulatory potential through enhancer-promoter chromatin interactions. SNP rs11631591 showed T-cell-specific enhancer activity and an allele-specific interaction with hnRNP-K protein. Inhibition of hnRNP-K protein by 5-FU decreased expression of *RASGRP1* in T-cells, suggesting that hnRNP-K plays an important role in *RASGRP1* expression through interactions with the risk genotype of SNP rs11631591. These results are consistent with this SNP being an important factor contributing to SLE pathogenicity.

Heterogeneous nuclear ribonucleoproteins (hnRNPs) represent a large family of nucleic acid-binding proteins implicated in various cellular processes including transcription and translation [24, 80]. hnRNP-K is a highly multifunctional protein, with annotated roles in chromatin remodeling, transcription, splicing and translation [80]. It is primarily referred to as an RNA-binding protein specific for “poly-C” repeats [81], but it actually prefers single-stranded DNA and can bind to double-stranded DNA [82]. hnRNP-K can act as a transcriptional activator or repressor [83]; notable examples include transcriptional repression of *CD43* in leukocytes [84] and transcriptional activation of *c-myc* in B-cells [85]. Its DNA-binding preference is found to be repeats of the CT motif, separated by several base pairs [82], confirmed by structure determination [86]. There are several CT motifs in the immediate environment of rs11631591, whose hnRNP-K binding could be affected by the SNP. It should also be noted that several of the other abundant proteins pulled down by the double-stranded DNA EMSA are primarily annotated as RNA-binding proteins, including hnRNP-M and splicing factor U2AF. Other transcription factors were also abundant, including far upstream element-binding protein 3, supporting the notion that this locus is indeed transcriptionally active.

Taken together, we have identified and mechanistically dissected a lupus risk locus in the 2^nd^ intron of *RASGRP1*, which regulates T- and B-cell development and the MAP kinase pathway. Single SNPs were found to control transcriptional activation and binding to several proteins, including the transcription factor hnRNP-K. Experiments confirmed that both the single base-pair risk-to-non-risk substitutions and pharmacological inhibition of hnRNP-K decreased MAPK signaling in T-cells. Systematic refinement of large GWAS peaks to single SNPs, combined with experimental mechanistic analysis, is critical to understand the genetics of highly multigenic diseases and to drive therapeutic interventions to improve human health.

## Acknowledgments

The authors thank all of the lupus patients and unaffected controls who participated in this study. We thank the research assistants, coordinators and physicians who helped in the recruitment of subjects, including the individuals in the coordinating projects.

## Web resources

Bentham and Morris summary SLE GWAS: http://insidegen.com/

---

**Supplementary Figure 1.**
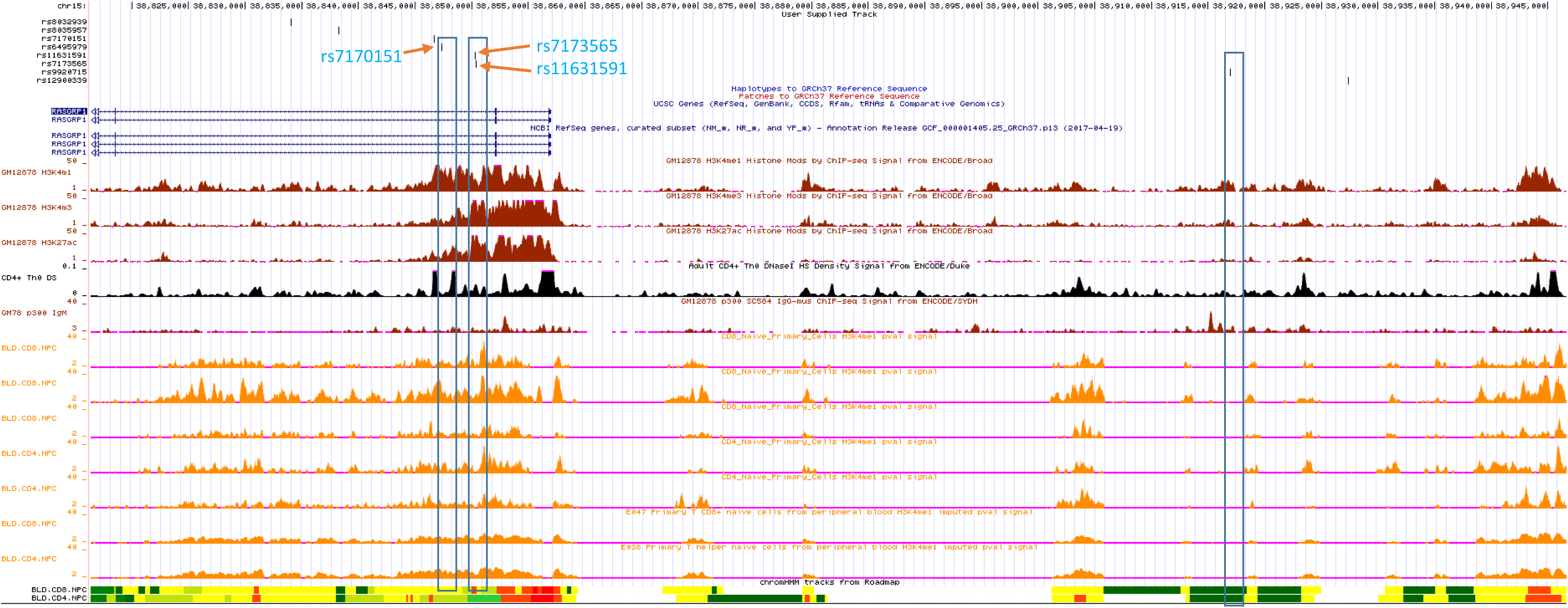
Collocation of our selected potentially functional SNPs with H2K27ac, H3K4me1, H3K4me3 and DNAse hypersensitivity in lymphoblastoid GM12878, Naïve CD4+, CD8+ T-cells and T-helper cells.

**Supplementary Figure 2.**
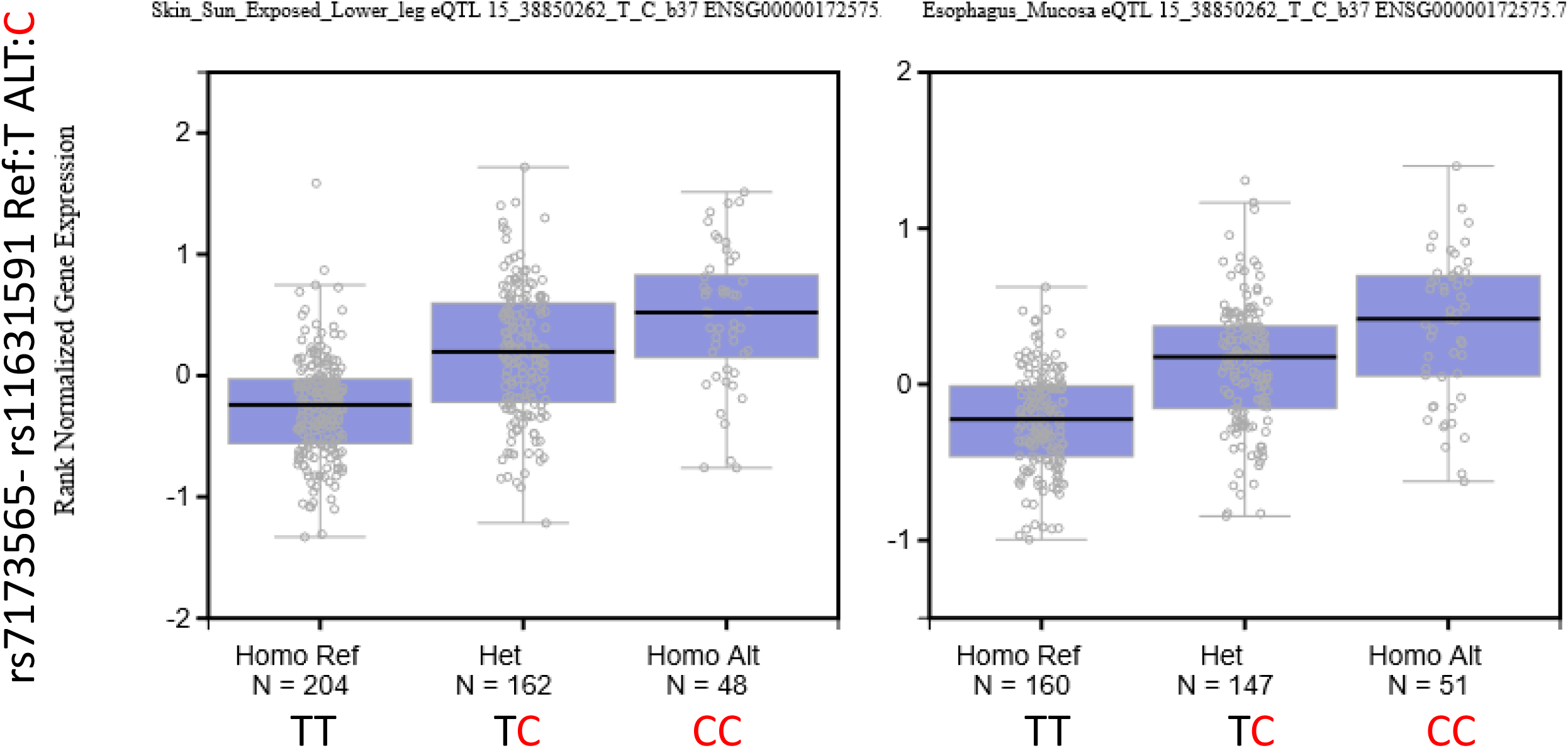
*RASGRP1* expression levels in multiple tissues (skin, left; esophagus, right) correlates with rs7173565-rs11631591 genotypes. Reference allele T; alternate allele C **(red).**

**Supplementary Figure 3.**
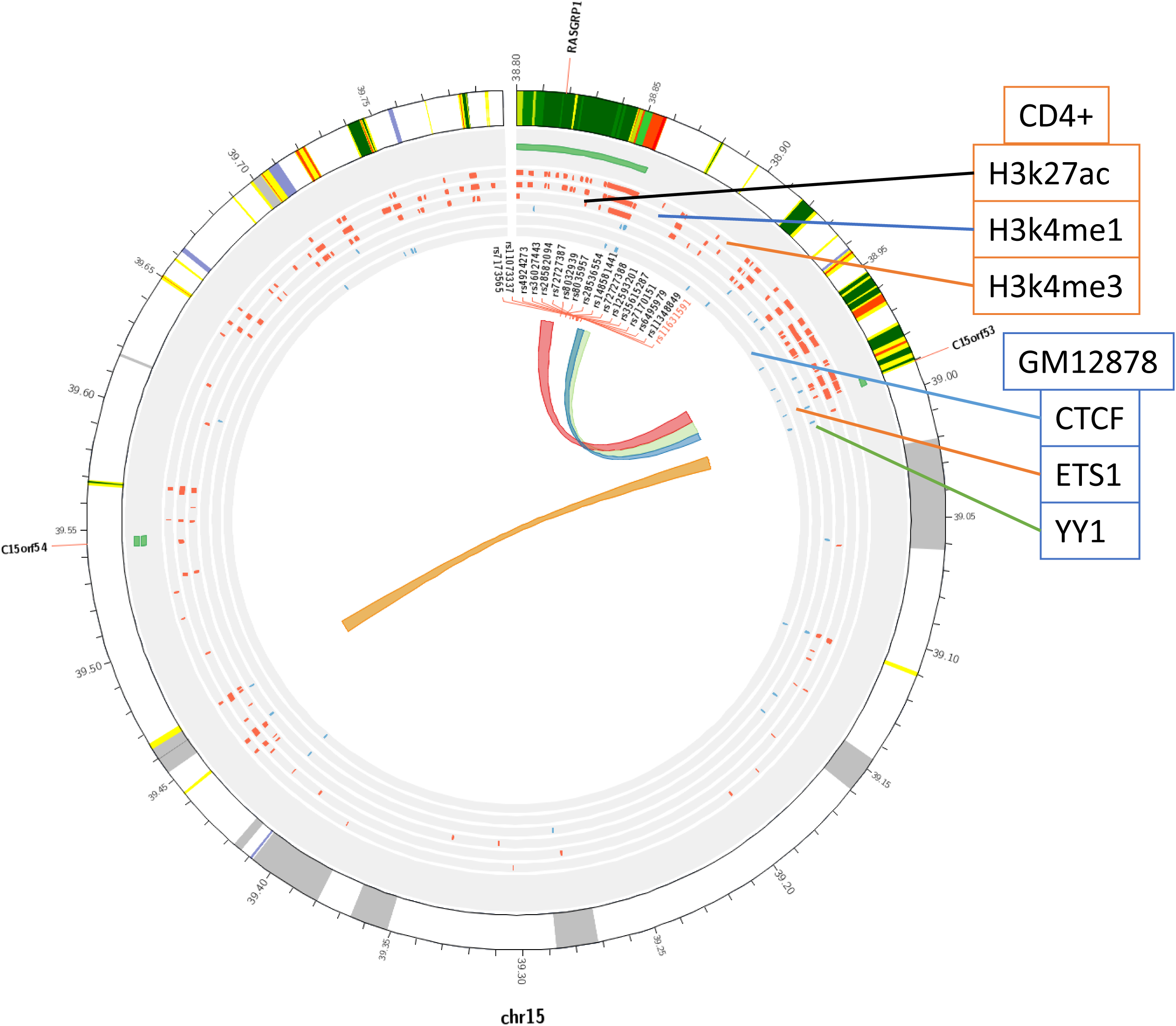
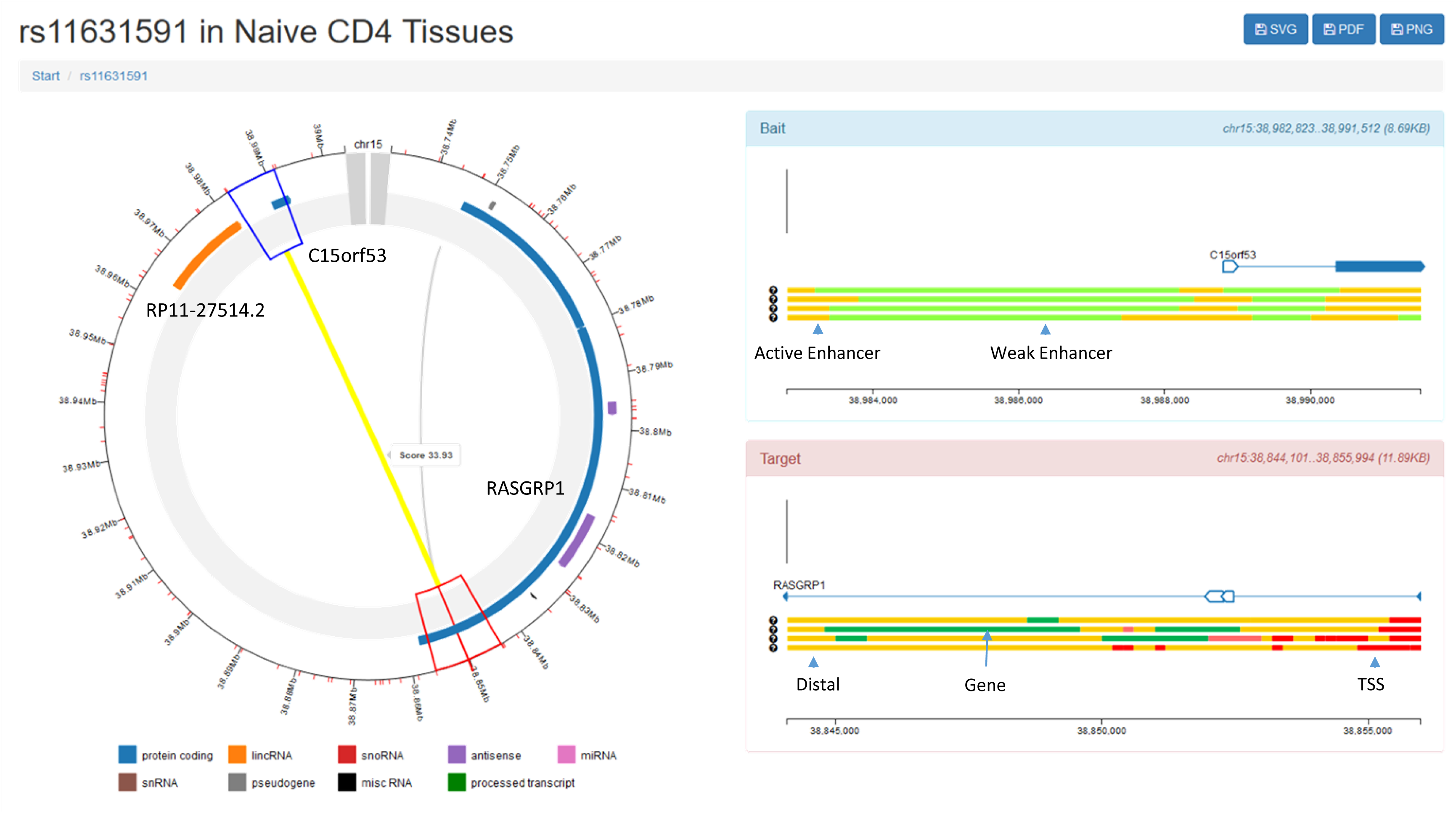
Chromatin interactions at the *RASGRP1* region overlap with rs11631591 in GM12878 cells. A. The three rings of orange marks show peaks for CD4^+^ T-cell H3K27ac, H3K4me1 and H3K4me3; the three rings of blue marks show peaks of CTCT, ETS1 and YY1 transcription factor binding. Colors in outer ring represent the 15-state classification of ChromHMM, notably: dark green for transcription, red for transcription start site/promoter, orange for enhancer. B. Promoter-capture Hi-C in Naïve CD4^+^ T-cells. The blue side (Bait) represents the promoter of *C15orf53* baited for the experiment. The red side (Target) represents the interaction side of the experiment. Intronic SNP rs11631591 lies in the target (*i.e.* enhancer) side of the interaction with the baited promoter of *C15orf53*. Bait and Target colors under the gene show chromatin state in CD4^+^ T-cells using the chromHMM 15-state classification.

**Supplementary Figure 4.**
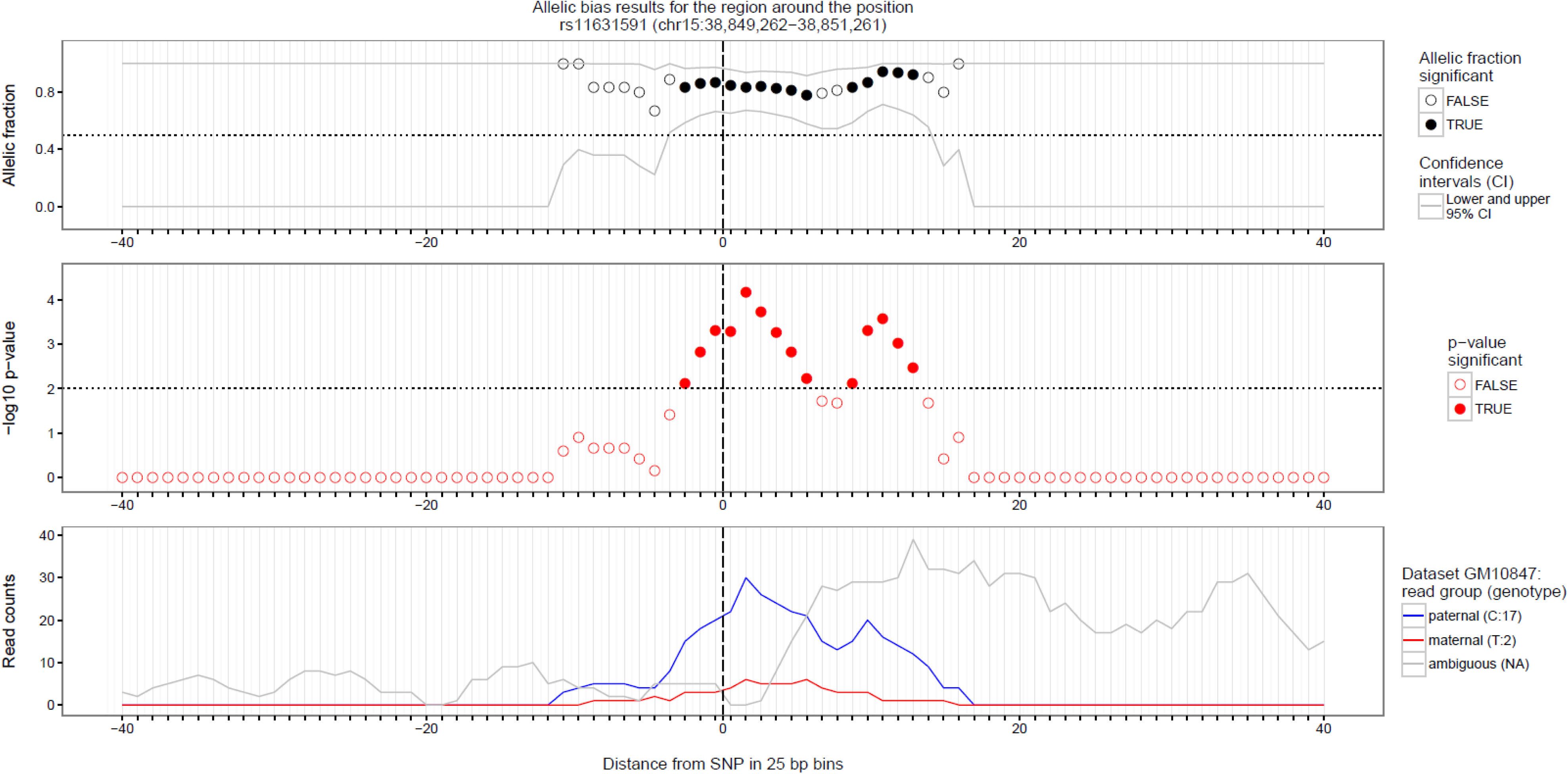
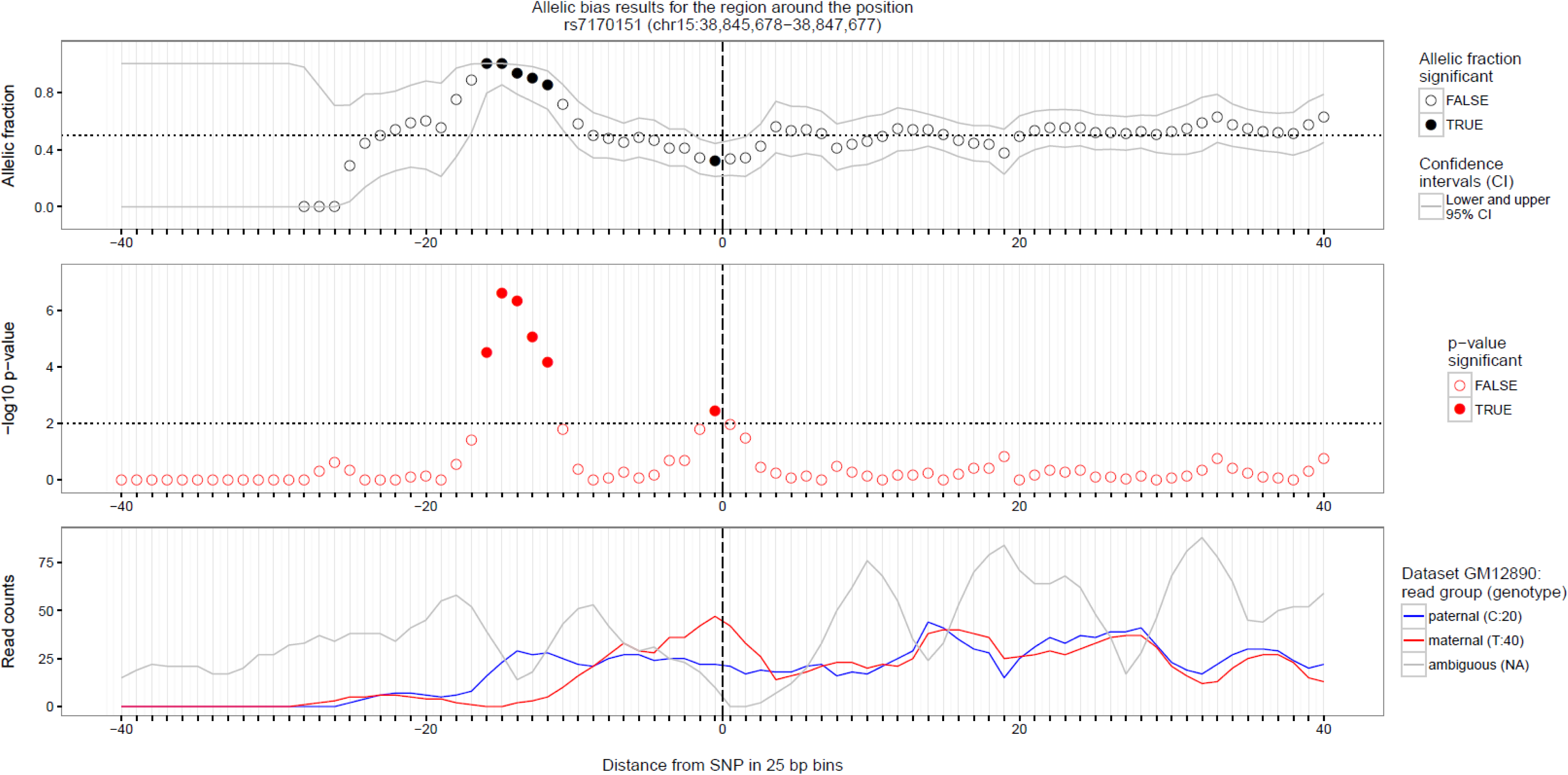
Allele-dependent binding to histones. A. H3K4me3 (rs11631591-rs7173565), and B. H3K4me1 (rs7170151) for lymphoblastoid cells as estimated with SNPhood (R-package). Top panel depicts the significant allelic fraction at each position relative to the SNP (position 0). Middle panel represents how significant the bias is at each position. Bottom panel shows the count of allele reads found in paternal (blue) and maternal (red) strands.

**Supplementary Figure 5.**
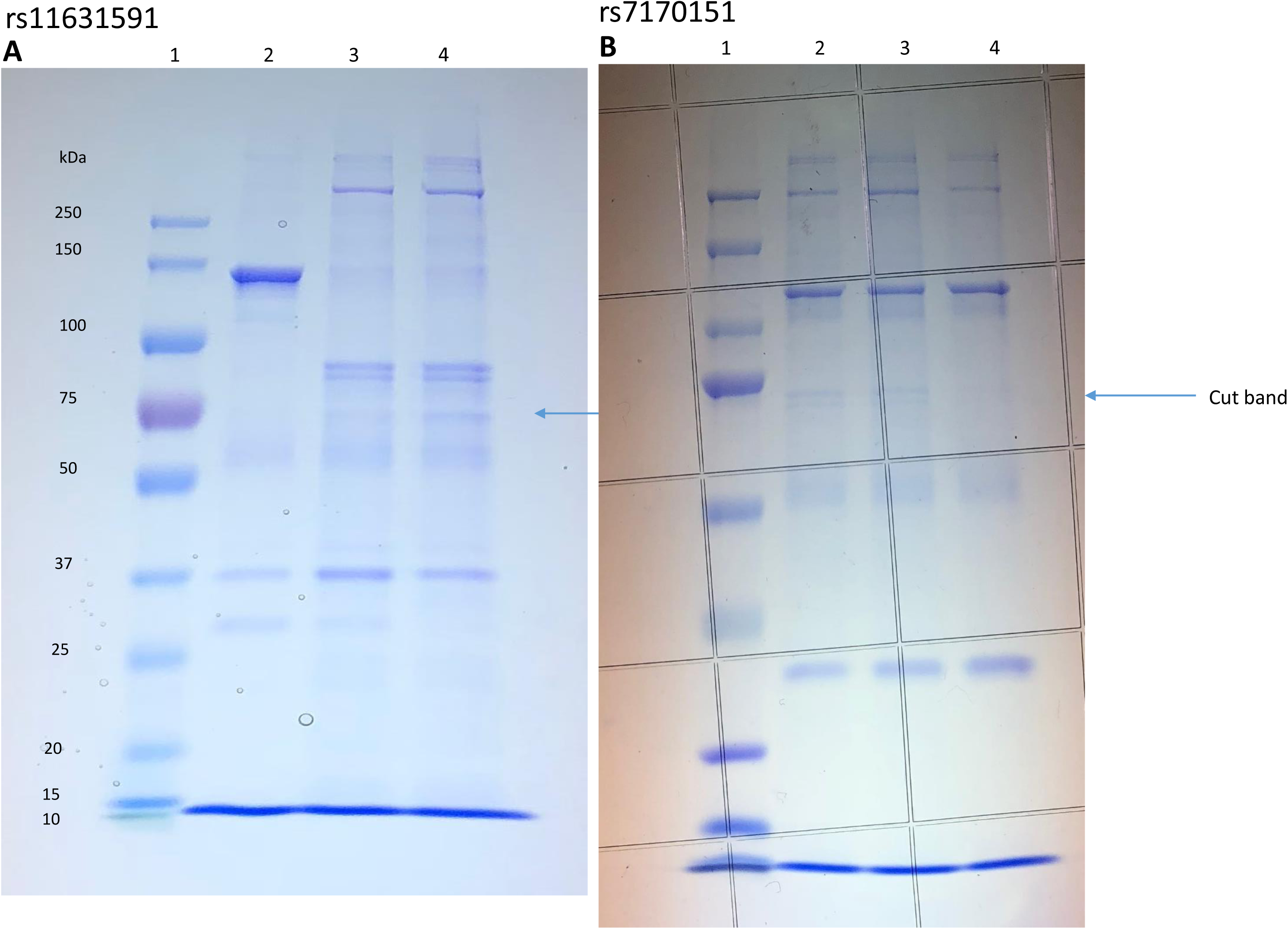
EMSA pulldown assay of rs11631591 (left) and rs7173565 (right) in Jurkat cells. EMSA was performed with biotin-labeled nucleotides containing the non-risk (TT) (41 bp) or risk (CC) (41 bp) polymorphisms along with the 5 bases-deleted SNP region (36 bp) for rs11631591 (**Figure 5A**). Nuclear extracts were derived from Jurkat cell lines. The ∼70 kDa apparent-MW protein/DNA complex shows allele-specific binding, which is absent in the deleted nucleotides. Similar results were also observed for the SNP (rs7170151) (**Figure 5B**). Lane 1: molecular weight marker; lane 2: nucleotide with deleted SNP region; lane 3: non-risk genotype; lane 4: risk genotype. Arrow shows the band of interest cut out for analysis by mass spectrometry.

**Supplementary Figure 6.**
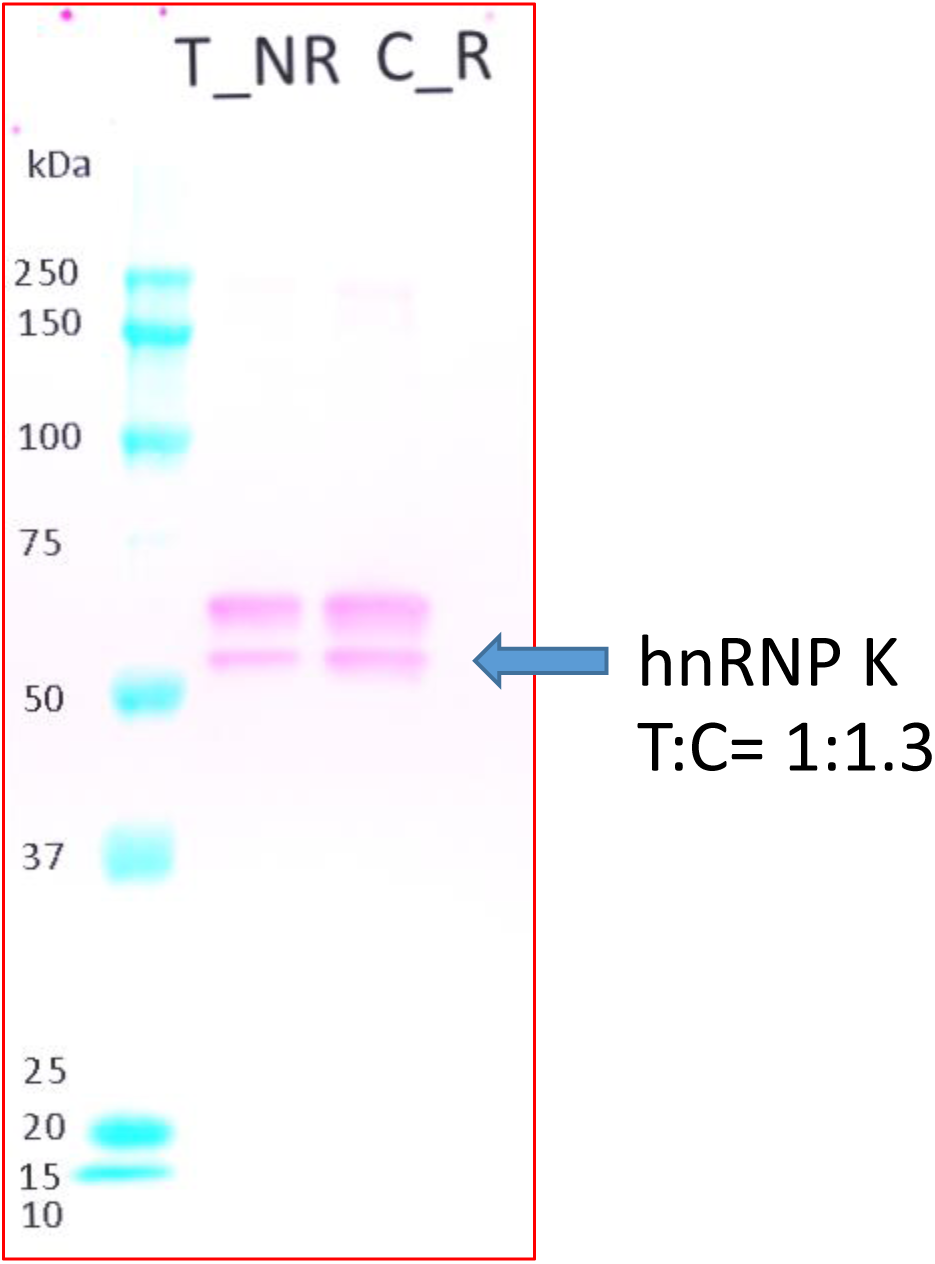
Western blot showing allele-specific binding of hnRNP-K against Jurkat cell lysate. The risk allele (C) of rs11631591 shows more binding than the non-risk allele (T). Densitometric measure of the bands T:C = 1:1.3.

**Supplementary Figure 7.**
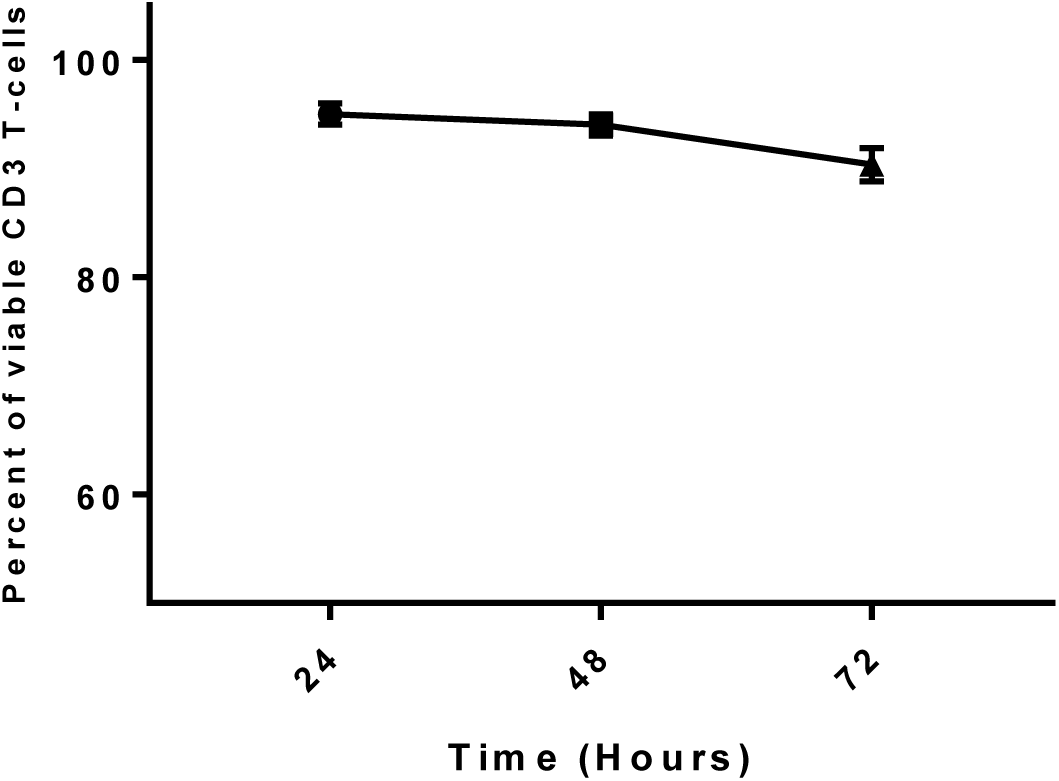
CD3^+^ T-cell viability after PMA stimulation. Cell viability was assessed by the trypan blue exclusion test, and results are expressed as the percentage of viable cells at each time point. Results represent the mean and SD of triplicate samples.

**Supplementary Figure 8.**
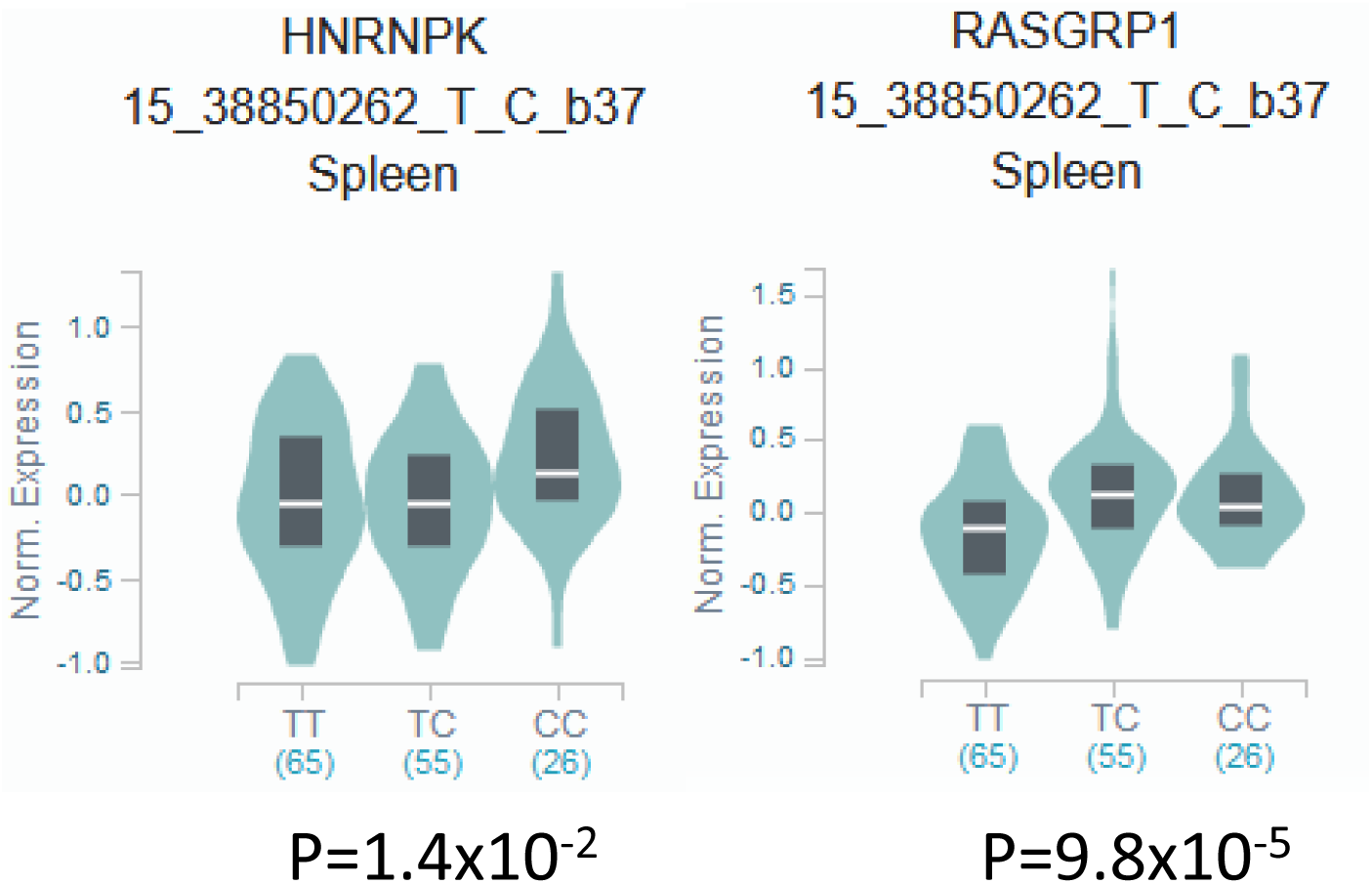
Correlation of the genotype of rs1163159 with *RASGRP1* and *hnRNP-K* in spleen.

**Supplementary Figure 9.**
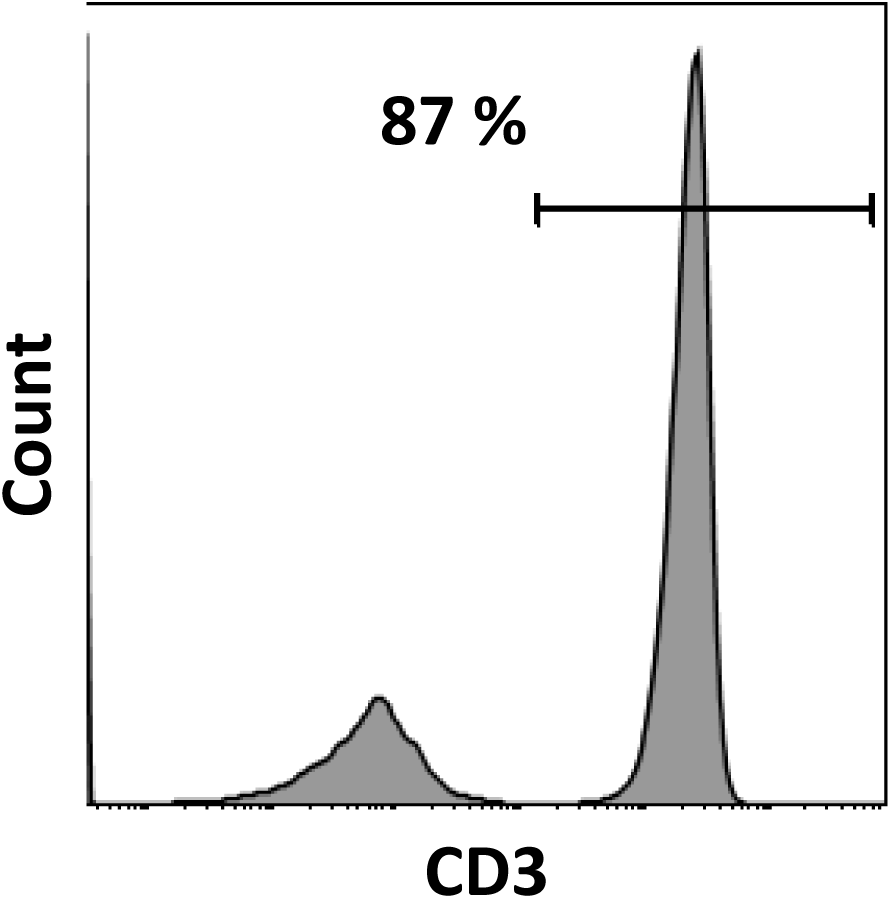
A representative flow cytometric analysis of CD3^+^ T cell population. CD3^+^ T cells were collected from PBMCs using MojoSort™ Human CD3^+^ T Cell Isolation Kit and were stained with fluorochrome-labeled antibodies CD3-PE and analyzed by flow cytometry. Histograms reflect the purity of isolated cells was 87 %.

**Supplementary Table 1.** Meta-analysis of all SNPs in our original report (supplementary table 5 from [4]). OR: odds ratio. 95% CI: 95% confidence interval. HetPVal: P-value for heterogeneity of association. Direction of OR: + for OR>1, -for OR<1.

**Supplementary Table 2.** Detailed annotation and prioritization for each genome-wide significant SNP. We integrated scores from 3dSNP, RegulomeDB and rSNP together with blood cell-specific information in eQTL, enhancer/super-enhancer existence, promoter capture HI-C (PCHiC), transcription factor binding site (TFBS) disruption and allele-specific expression/binding (ASE/ASB) into a weighted score for SNP prioritization. We chose the top three SNPs for further validation (red; rs11631591 and rs7173565 were used together because of the short distance between them).

**Supplementary Table 3.** Sequences of oligonucleotides used in EMSA and qPCR.

**Supplementary Table 4.** eQTLs identified for all candidate SNPs from GTeX, BloodeQTL, Geuvadis, Blueprint, and the Bios single-cell RNA project.

**Supplementary Table 5.** Methylation QTL. Methylation QTL assessed in immune cell lines from the BLUEPRINT project. FDR: False discovery rate.

**Supplementary Table 6.** Enhancer and super-enhancer location relative to all candidate SNPs.

**Supplementary Table 7.** Promoter capture Hi-C location for 14 cell lines.

**Supplementary Table 8.** Quantitative cytokine assessment for all candidate SNPs.

**Supplementary Table 9.** Allele-dependent expression and binding of H2K27ac and H3K4me1 from the BLUEPRINT for the candidate SNPs.

**Supplementary Table 10.** Allele-dependent transcription factor binding site (TFBS) disruption for all candidate SNPs from HT-Selex, Hocomoco, SwissRegulon, HOMER and JASPER databases.

**Supplementary Table 11.** Protein capture identification for rs11631591 risk *vs.* non-risk allele in Jurkat cells.

**Supplementary Table 12.** Characteristics of CD3+ T-cell samples used in this study.

## Literature Cited

1. Alarcon GS, McGwin G, Jr., Petri M, Reveille JD, Ramsey-Goldman R, Kimberly RP, Group PS: Baseline characteristics of a multiethnic lupus cohort: PROFILE. Lupus 2002, 11:95–101.

2. Molineros JE, Yang W, Zhou XJ, Sun C, Okada Y, Zhang H, Heng Chua K, Lau YL, Kochi Y, Suzuki A, et al: Confirmation of five novel susceptibility loci for systemic lupus erythematosus (SLE) and integrated network analysis of 82 SLE susceptibility loci. Hum Mol Genet 2017, 26:1205–1216.

3. Langefeld CD, Ainsworth HC, Cunninghame Graham DS, Kelly JA, Comeau ME, Marion MC, Howard TD, Ramos PS, Croker JA, Morris DL, et al: Transancestral mapping and genetic load in systemic lupus erythematosus. Nat Commun 2017, 8:16021.

4. Sun C, Molineros JE, Looger LL, Zhou XJ, Kim K, Okada Y, Ma J, Qi YY, Kim-Howard X, Motghare P, et al: High-density genotyping of immune-related loci identifies new SLE risk variants in individuals with Asian ancestry. Nat Genet 2016.

5. Carrasco S, Merida I: Diacylglycerol-dependent binding recruits PKCtheta and RasGRP1 C1 domains to specific subcellular localizations in living T lymphocytes. Mol Biol Cell 2004, 15:2932–2942.

6. Beaulieu N, Zahedi B, Goulding RE, Tazmini G, Anthony KV, Omeis SL, de Jong DR, Kay RJ: Regulation of RasGRP1 by B cell antigen receptor requires cooperativity between three domains controlling translocation to the plasma membrane. Mol Biol Cell 2007, 18:3156–3168.

7. Ebinu JO, Bottorff DA, Chan EY, Stang SL, Dunn RJ, Stone JC: RasGRP, a Ras guanyl nucleotide-releasing protein with calcium- and diacylglycerol-binding motifs. Science 1998, 280:1082– 1086.

8. Dower NA, Stang SL, Bottorff DA, Ebinu JO, Dickie P, Ostergaard HL, Stone JC: RasGRP is essential for mouse thymocyte differentiation and TCR signaling. Nat Immunol 2000, 1:317– 321.

9. Liu Y, Zhu M, Nishida K, Hirano T, Zhang W: An essential role for RasGRP1 in mast cell function and IgE-mediated allergic response. J Exp Med 2007, 204:93–103.

10. Coughlin JJ, Stang SL, Dower NA, Stone JC: RasGRP1 and RasGRP3 regulate B cell proliferation by facilitating B cell receptor-Ras signaling. J Immunol 2005, 175:7179–7184.

11. Shen S, Chen Y, Gorentla BK, Lu J, Stone JC, Zhong XP: Critical roles of RasGRP1 for invariant NKT cell development. J Immunol 2011, 187:4467–4473.

12. Bartlett A, Buhlmann JE, Stone J, Lim B, Barrington RA: Multiple checkpoint breach of B cell tolerance in Rasgrp1-deficient mice. J Immunol 2013, 191:3605–3613.

13. Guo B, Rothstein TL: RasGRP1 Is an Essential Signaling Molecule for Development of B1a Cells with Autoantigen Receptors. J Immunol 2016, 196:2583–2590.

14. Priatel JJ, Teh SJ, Dower NA, Stone JC, Teh HS: RasGRP1 transduces low-grade TCR signals which are critical for T cell development, homeostasis, and differentiation. Immunity 2002, 17:617–627.

15. Salzer E, Cagdas D, Hons M, Mace EM, Garncarz W, Petronczki OY, Platzer R, Pfajfer L, Bilic I, Ban SA, et al: RASGRP1 deficiency causes immunodeficiency with impaired cytoskeletal dynamics. Nat Immunol 2016, 17:1352–1360.

16. Winter S, Martin E, Boutboul D, Lenoir C, Boudjemaa S, Petit A, Picard C, Fischer A, Leverger G, Latour S: Loss of RASGRP1 in humans impairs T-cell expansion leading to Epstein-Barr virus susceptibility. EMBO Mol Med 2018, 10:188–199.

17. Mao H, Yang W, Latour S, Yang J, Winter S, Zheng J, Ni K, Lv M, Liu C, Huang H, et al: RASGRP1 mutation in autoimmune lymphoproliferative syndrome-like disease. J Allergy Clin Immunol 2018, 142:595–604 e516.

18. Golinski ML, Vandhuick T, Derambure C, Freret M, Lecuyer M, Guillou C, Hiron M, Boyer O, Le Loet X, Vittecoq O, Lequerre T: Dysregulation of RasGRP1 in rheumatoid arthritis and modulation of RasGRP3 as a biomarker of TNFalpha inhibitors. Arthritis Res Ther 2015, 17:382.

19. Rapoport MJ, Bloch O, Amit-Vasina M, Yona E, Molad Y: Constitutive abnormal expression of RasGRP-1 isoforms and low expression of PARP-1 in patients with systemic lupus erythematosus. Lupus 2011, 20:1501–1509.

20. Yasuda S, Stevens RL, Terada T, Takeda M, Hashimoto T, Fukae J, Horita T, Kataoka H, Atsumi T, Koike T: Defective expression of Ras guanyl nucleotide-releasing protein 1 in a subset of patients with systemic lupus erythematosus. J Immunol 2007, 179:4890–4900.

21. Kono M, Kurita T, Yasuda S, Kono M, Fujieda Y, Bohgaki T, Katsuyama T, Tsokos GC, Moulton VR, Atsumi T: Decreased expression of Serine/arginine-rich splicing factor 1 in T cells from patients with active systemic lupus erythematosus accounts for reduced expression of RasGRP1 and DNA methyltransferase 1. Arthritis Rheumatol 2018.

22. Deng C, Lu Q, Zhang Z, Rao T, Attwood J, Yung R, Richardson B: Hydralazine may induce autoimmunity by inhibiting extracellular signal-regulated kinase pathway signaling. Arthritis Rheum 2003, 48:746–756.

23. Molad Y, Amit-Vasina M, Bloch O, Yona E, Rapoport MJ: Increased ERK and JNK activities correlate with disease activity in patients with systemic lupus erythematosus. Ann Rheum Dis 2010, 69:175–180.

24. Bloch O, Amit-Vazina M, Yona E, Molad Y, Rapoport MJ: Increased ERK and JNK activation and decreased ERK/JNK ratio are associated with long-term organ damage in patients with systemic lupus erythematosus. Rheumatology (Oxford) 2014, 53:1034–1042.

25. Rapoport MJ, Amit M, Aharoni D, Weiss M, Weissgarten J, Bruck N, Buchs A, Bistritzer T, Molad Y: Constitutive up-regulated activity of MAP kinase is associated with down-regulated early p21Ras pathway in lymphocytes of SLE patients. J Autoimmun 2002, 19:63–70.

26. Bentham J, Morris DL, Cunninghame Graham DS, Pinder CL, Tombleson P, Behrens TW, Martin J, Fairfax BP, Knight JC, Chen L, et al: Genetic association analyses implicate aberrant regulation of innate and adaptive immunity genes in the pathogenesis of systemic lupus erythematosus. Nat Genet 2015, 47:1457–1464.

27. Morris DL, Sheng Y, Zhang Y, Wang YF, Zhu Z, Tombleson P, Chen L, Cunninghame Graham DS, Bentham J, Roberts AL, et al: Genome-wide association meta-analysis in Chinese and European individuals identifies ten new loci associated with systemic lupus erythematosus. Nat Genet 2016.

28. Okada Y, Shimane K, Kochi Y, Tahira T, Suzuki A, Higasa K, Takahashi A, Horita T, Atsumi T, Ishii T, et al: A genome-wide association study identified AFF1 as a susceptibility locus for systemic lupus eyrthematosus in Japanese. PLoS Genet 2012, 8:e1002455.

29. Hochberg MC: Updating the American College of Rheumatology revised criteria for the classification of systemic lupus erythematosus. Arthritis Rheum 1997, 40:1725.

30. Purcell S, Neale B, Todd-Brown K, Thomas L, Ferreira MA, Bender D, Maller J, Sklar P, de Bakker PI, Daly MJ, Sham PC: PLINK: a tool set for whole-genome association and population-based linkage analyses. Am J Hum Genet 2007, 81:559–575.

31. Willer CJ, Li Y, Abecasis GR: METAL: fast and efficient meta-analysis of genomewide association scans. Bioinformatics 2010, 26:2190–2191.

32. Ward LD, Kellis M: HaploReg: a resource for exploring chromatin states, conservation, and regulatory motif alterations within sets of genetically linked variants. Nucleic Acids Res 2012, 40:D930–934.

33. Consortium EP: An integrated encyclopedia of DNA elements in the human genome. Nature 2012, 489:57–74.

34. Pradel LC, Vanhille L, Spicuglia S: [The European Blueprint project: towards a full epigenome characterization of the immune system]. Med Sci (Paris) 2015, 31:236–238.

35. Lu Y, Quan C, Chen H, Bo X, Zhang C: 3DSNP: a database for linking human noncoding SNPs to their three-dimensional interacting genes. Nucleic Acids Res 2017, 45:D643–D649.

36. Boyle AP, Hong EL, Hariharan M, Cheng Y, Schaub MA, Kasowski M, Karczewski KJ, Park J, Hitz BC, Weng S, et al: Annotation of functional variation in personal genomes using RegulomeDB. Genome Res 2012, 22:1790–1797.

37. Guo L, Du Y, Chang S, Zhang K, Wang J: rSNPBase: a database for curated regulatory SNPs. Nucleic Acids Res 2014, 42:D1033–1039.

38. Lappalainen T, Sammeth M, Friedlander MR, Hoen PA, Monlong J, Rivas MA, Gonzalez-Porta M, Kurbatova N, Griebel T, Ferreira PG, et al: Transcriptome and genome sequencing uncovers functional variation in humans. Nature 2013, 501:506–511.

39. Westra HJ, Peters MJ, Esko T, Yaghootkar H, Schurmann C, Kettunen J, Christiansen MW, Fairfax BP, Schramm K, Powell JE, et al: Systematic identification of trans eQTLs as putative drivers of known disease associations. Nat Genet 2013, 45:1238–1243.

40. van der Wijst MGP, Brugge H, de Vries DH, Deelen P, Swertz MA, LifeLines Cohort S, Consortium B, Franke L: Single-cell RNA sequencing identifies celltype-specific cis-eQTLs and co-expression QTLs. Nat Genet 2018, 50:493–497.

41. Consortium GT: Human genomics. The Genotype-Tissue Expression (GTEx) pilot analysis: multitissue gene regulation in humans. Science 2015, 348:648–660.

42. Chen L, Ge B, Casale FP, Vasquez L, Kwan T, Garrido-Martin D, Watt S, Yan Y, Kundu K, Ecker S, et al: Genetic Drivers of Epigenetic and Transcriptional Variation in Human Immune Cells. Cell 2016, 167:1398–1414 e1324.

43. Coetzee SG, Coetzee GA, Hazelett DJ: motifbreakR: an R/Bioconductor package for predicting variant effects at transcription factor binding sites. Bioinformatics 2015, 31:3847–3849.

44. Khan A, Zhang X: dbSUPER: a database of super-enhancers in mouse and human genome. Nucleic Acids Res 2016, 44:D164–171.

45. Corradin O, Saiakhova A, Akhtar-Zaidi B, Myeroff L, Willis J, Cowper-Sal lari R, Lupien M, Markowitz S, Scacheri PC: Combinatorial effects of multiple enhancer variants in linkage disequilibrium dictate levels of gene expression to confer susceptibility to common traits. Genome Res 2014, 24:1–13.

46. Gao T, He B, Liu S, Zhu H, Tan K, Qian J: EnhancerAtlas: a resource for enhancer annotation and analysis in 105 human cell/tissue types. Bioinformatics 2016.

47. Li R, Liu Y, Li T, Li C: 3Disease Browser: A Web server for integrating 3D genome and disease-associated chromosome rearrangement data. Sci Rep 2016, 6:34651.

48. Schofield EC, Carver T, Achuthan P, Freire-Pritchett P, Spivakov M, Todd JA, Burren OS: CHiCP: a web-based tool for the integrative and interactive visualization of promoter capture Hi-C datasets. Bioinformatics 2016, 32:2511–2513.

49. Rao SS, Huntley MH, Durand NC, Stamenova EK, Bochkov ID, Robinson JT, Sanborn AL, Machol I, Omer AD, Lander ES, Aiden EL: A 3D map of the human genome at kilobase resolution reveals principles of chromatin looping. Cell 2014, 159:1665–1680.

50. Sanborn AL, Rao SS, Huang SC, Durand NC, Huntley MH, Jewett AI, Bochkov ID, Chinnappan D, Cutkosky A, Li J, et al: Chromatin extrusion explains key features of loop and domain formation in wild-type and engineered genomes. Proc Natl Acad Sci U S A 2015, 112:E6456–6465.

51. Cairns J, Freire-Pritchett P, Wingett SW, Varnai C, Dimond A, Plagnol V, Zerbino D, Schoenfelder S, Javierre BM, Osborne C, et al: CHiCAGO: robust detection of DNA looping interactions in Capture Hi-C data. Genome Biol 2016, 17:127.

52. Taberlay PC, Achinger-Kawecka J, Lun AT, Buske FA, Sabir K, Gould CM, Zotenko E, Bert SA, Giles KA, Bauer DC, et al: Three-dimensional disorganization of the cancer genome occurs coincident with long-range genetic and epigenetic alterations. Genome Res 2016, 26:719–73x1.

53. Arnold C, Bhat P, Zaugg JB: SNPhood: investigate, quantify and visualise the epigenomic neighbourhood of SNPs using NGS data. Bioinformatics 2016, 32:2359–2360.

54. Xiao Z, Ko HL, Goh EH, Wang B, Ren EC: hnRNP K suppresses apoptosis independent of p53 status by maintaining high levels of endogenous caspase inhibitors. Carcinogenesis 2013, 34:1458–1467.

55. Wang JC, Foroud T, Hinrichs AL, Le NX, Bertelsen S, Budde JP, Harari O, Koller DL, Wetherill L, Agrawal A, et al: A genome-wide association study of alcohol-dependence symptom counts in extended pedigrees identifies C15orf53. Mol Psychiatry 2013, 18:1218–1224.

56. Okada Y, Wu D, Trynka G, Raj T, Terao C, Ikari K, Kochi Y, Ohmura K, Suzuki A, Yoshida S, et al: Genetics of rheumatoid arthritis contributes to biology and drug discovery. Nature 2014, 506:376–381.

57. Grant SF, Qu HQ, Bradfield JP, Marchand L, Kim CE, Glessner JT, Grabs R, Taback SP, Frackelton EC, Eckert AW, et al: Follow-up analysis of genome-wide association data identifies novel loci for type 1 diabetes. Diabetes 2009, 58:290–295.

58. Hnisz D, Abraham BJ, Lee TI, Lau A, Saint-Andre V, Sigova AA, Hoke HA, Young RA: Super-enhancers in the control of cell identity and disease. Cell 2013, 155:934–947.

59. Davis LS, Hutcheson J, Mohan C: The role of cytokines in the pathogenesis and treatment of systemic lupus erythematosus. J Interferon Cytokine Res 2011, 31:781–789.

60. Han JW, Zheng HF, Cui Y, Sun LD, Ye DQ, Hu Z, Xu JH, Cai ZM, Huang W, Zhao GP, et al: Genome-wide association study in a Chinese Han population identifies nine new susceptibility loci for systemic lupus erythematosus. Nat Genet 2009, 41:1234–1237.

61. International Consortium for Systemic Lupus Erythematosus G, Harley JB, Alarcon-Riquelme ME, Criswell LA, Jacob CO, Kimberly RP, Moser KL, Tsao BP, Vyse TJ, Langefeld CD, et al: Genome-wide association scan in women with systemic lupus erythematosus identifies susceptibility variants in ITGAM, PXK, KIAA1542 and other loci. Nat Genet 2008, 40:204–210.

62. Javierre BM, Burren OS, Wilder SP, Kreuzhuber R, Hill SM, Sewitz S, Cairns J, Wingett SW, Varnai C, Thiecke MJ, et al: Lineage-Specific Genome Architecture Links Enhancers and Non-coding Disease Variants to Target Gene Promoters. Cell 2016, 167:1369–1384 e1319.

63. Cairns BR: The logic of chromatin architecture and remodelling at promoters. Nature 2009, 461:193–198.

64. Tillo D, Kaplan N, Moore IK, Fondufe-Mittendorf Y, Gossett AJ, Field Y, Lieb JD, Widom J, Segal E, Hughes TR: High nucleosome occupancy is encoded at human regulatory sequences. PLoS One 2010, 5:e9129.

65. Wiench M, John S, Baek S, Johnson TA, Sung MH, Escobar T, Simmons CA, Pearce KH, Biddie SC, Sabo PJ, et al: DNA methylation status predicts cell type-specific enhancer activity. EMBO J 2011, 30:3028–3039.

66. Trynka G, Sandor C, Han B, Xu H, Stranger BE, Liu XS, Raychaudhuri S: Chromatin marks identify critical cell types for fine mapping complex trait variants. Nat Genet 2013, 45:124–130.

67. Pasquali L, Gaulton KJ, Rodriguez-Segui SA, Mularoni L, Miguel-Escalada I, Akerman I, Tena JJ, Moran I, Gomez-Marin C, van de Bunt M, et al: Pancreatic islet enhancer clusters enriched in type 2 diabetes risk-associated variants. Nat Genet 2014, 46:136–143.

68. Raj T, Rothamel K, Mostafavi S, Ye C, Lee MN, Replogle JM, Feng T, Lee M, Asinovski N, Frohlich I, et al: Polarization of the effects of autoimmune and neurodegenerative risk alleles in leukocytes. Science 2014, 344:519–523.

69. Heinz S, Romanoski CE, Benner C, Glass CK: The selection and function of cell type-specific enhancers. Nat Rev Mol Cell Biol 2015, 16:144–154.

70. Degner JF, Pai AA, Pique-Regi R, Veyrieras JB, Gaffney DJ, Pickrell JK, De Leon S, Michelini K, Lewellen N, Crawford GE, et al: DNase I sensitivity QTLs are a major determinant of human expression variation. Nature 2012, 482:390–394.

71. Kasowski M, Kyriazopoulou-Panagiotopoulou S, Grubert F, Zaugg JB, Kundaje A, Liu Y, Boyle AP, Zhang QC, Zakharia F, Spacek DV, et al: Extensive variation in chromatin states across humans. Science 2013, 342:750–752.

72. Kilpinen H, Waszak SM, Gschwind AR, Raghav SK, Witwicki RM, Orioli A, Migliavacca E, Wiederkehr M, Gutierrez-Arcelus M, Panousis NI, et al: Coordinated effects of sequence variation on DNA binding, chromatin structure, and transcription. Science 2013, 342:744–747.

73. McVicker G, van de Geijn B, Degner JF, Cain CE, Banovich NE, Raj A, Lewellen N, Myrthil M, Gilad Y, Pritchard JK: Identification of genetic variants that affect histone modifications in human cells. Science 2013, 342:747–749.

74. Gaffney DJ, Veyrieras JB, Degner JF, Pique-Regi R, Pai AA, Crawford GE, Stephens M, Gilad Y, Pritchard JK: Dissecting the regulatory architecture of gene expression QTLs. Genome Biol 2012, 13:R7.

75. Helgadottir A, Thorleifsson G, Manolescu A, Gretarsdottir S, Blondal T, Jonasdottir A, Jonasdottir A, Sigurdsson A, Baker A, Palsson A, et al: A common variant on chromosome 9p21 affects the risk of myocardial infarction. Science 2007, 316:1491–1493.

76. Gaulton KJ, Nammo T, Pasquali L, Simon JM, Giresi PG, Fogarty MP, Panhuis TM, Mieczkowski P, Secchi A, Bosco D, et al: A map of open chromatin in human pancreatic islets. Nat Genet 2010, 42:255–259.

77. Kasowski M, Grubert F, Heffelfinger C, Hariharan M, Asabere A, Waszak SM, Habegger L, Rozowsky J, Shi M, Urban AE, et al: Variation in transcription factor binding among humans. Science 2010, 328:232–235.

78. Cowper-Sal lari R, Zhang X, Wright JB, Bailey SD, Cole MD, Eeckhoute J, Moore JH, Lupien M: Breast cancer risk-associated SNPs modulate the affinity of chromatin for FOXA1 and alter gene expression. Nat Genet 2012, 44:1191–1198.

79. Bauer DE, Kamran SC, Lessard S, Xu J, Fujiwara Y, Lin C, Shao Z, Canver MC, Smith EC, Pinello L, et al: An erythroid enhancer of BCL11A subject to genetic variation determines fetal hemoglobin level. Science 2013, 342:253–257.

80. Bomsztyk K, Denisenko O, Ostrowski J: hnRNP K: one protein multiple processes. Bioessays 2004, 26:629–638.

81. Leffers H, Dejgaard K, Celis JE: Characterisation of two major cellular poly(rC)-binding human proteins, each containing three K-homologous (KH) domains. Eur J Biochem 1995, 230:447–453.

82. Tomonaga T, Levens D: Heterogeneous nuclear ribonucleoprotein K is a DNA-binding transactivator. J Biol Chem 1995, 270:4875–4881.

83. Choi HS, Hwang CK, Song KY, Law PY, Wei LN, Loh HH: Poly(C)-binding proteins as transcriptional regulators of gene expression. Biochem Biophys Res Commun 2009, 380:431– 436.

84. Da Silva N, Bharti A, Shelley CS: hnRNP-K and Pur(alpha) act together to repress the transcriptional activity of the CD43 gene promoter. Blood 2002, 100:3536–3544.

85. Michelotti GA, Michelotti EF, Pullner A, Duncan RC, Eick D, Levens D: Multiple single-stranded cis elements are associated with activated chromatin of the human c-myc gene in vivo. Mol Cell Biol 1996, 16:2656–2669.

86. Braddock DT, Baber JL, Levens D, Clore GM: Molecular basis of sequence-specific single-stranded DNA recognition by KH domains: solution structure of a complex between hnRNP K KH3 and single-stranded DNA. EMBO J 2002, 21:3476–3485.

